# Genetic Code Evolution Reconstructed with Aligned Metrics

**DOI:** 10.1101/2020.07.29.227728

**Authors:** Brian K. Davis

**Affiliations:** Research Foundation of Southern California Inc., 8861 Villa La Jolla Dr. #13595, La Jolla, CA 92037.

**Keywords:** path-length, depth, hydropathy, enthalpy states, fidelity, early proteins, codon bias, mutations

## Abstract

Sequence homology in pre-divergence tRNA species revealed cofactor/adaptors cognate for 16 amino acids derived from oxaloacetate, pyruvate, phosphoglycerate, or phosphoenolpyruvate were related. Synthesis path-distances of these amino acids correlated with phylogenetic depth, reflecting relative residue frequency in pre-divergence sequences. Both metrics were thus aligned in the four sub-families of the Aspartate family, and misaligned in the small Glutamate family; a functional difference was noted and seen to parallel synthetase duality. Amino acid synthetic order, based on path-distances, indicate NH_4_^+^ fixer amino acids, Asp^1^, Asn^2^, and homologues, Glu^1^, Gln^2^, formed the first code. Together with a termination signal, they acquired all four triplet 4-sets in the XAN column (X, 5’ coding site; N, any 3’-base). An invariant mid-A conformed with pre-code translation on a poly(A) template by a ratchet-equipped ribosome resulting in random, polyanionic polypeptides. Code expansion occurred in a compact (mutation minimizing) columnwise pattern, (XAN) ➔ XCN ➔ XGN ➔ XUN; with increasing mean path-distance, (1.5) ➔ 4 ➔ 5 ➔ 7 steps; amino acid side-chain hydrophobicity, (+6.6) ➔ −0.8 ➔ −1.5 ➔ −3.2 kcal/ mol; codon:anticodon H bond enthalpy (selection for bond-strength), (−12.5) ➔ −17.5 ➔ −15.5 ➔ −14.5 kcal/ mol; and precursor-specific 5’-base, A, oxaloacetate, G, pyruvate/oxaloacetate, U, phosphoglycerate/oxaloacetate, C, oxoglutarate, forming horizontal code domains. Codon bias evidence corroborated the XCN ➔ XGN step in expansion, and revealed row GNN coevolved with ANN, on correction for overprinting. Extended surface-attachment (Fajan-Paneth principle) by pro-Fd[5] and bilayer partitioning by H^+^ ATPase proteolipid-h1 subunit implicated expansion phase proteins in driving increases in side-chain hydrophobicity during code expansion. 3’-Base recruitment in pre-assigned codon boxes added six long (9-to 14-step) path amino acid, bearing a basic, or cyclic, side-chain; 3 of 4 polar, post-expansion amino acids acquired polar cluster NAN codons and 2 of 3 non-polar (Ile^7^ included) acquired non-polar cluster NUN codons, yieldng a split-box pair homology of *p* = 5.4×10^-3^. All eight overprinted codon boxes (GA^Y^_R_ for Asp^1^, Glu^1^ included) exhibit weak codon:anticodon H-bond enthalpy, −14 kcal/mol or higher, in three of six distinct code enthalpy states.

## 1. Introduction

Path-distances in the pre-LUCA reaction sequences that led to the standard set of amino acids in proteins (Crick, 1958) place both diacid amino acids, Asp^1^ and Glu^1^, and their amides, Asn^2^ and Gln^2^, at the origin of the genetic code (Davis, 1999).^1^ Together with a peptide chain-termination signal, they form a compact, error-minimizing set of four homologues, synthesized on paths of minimal length, and encoded by XAN triplets. The path-distance metric of amino acid synthetic order thus strategically positions the first code, in template-directed translation, at the point of entry of N atoms, contributed by ammonium ions, into the ancient pathways of amino acid synthesis. These paths originated at oxaloacetate, or 2-oxoglutarate, within the reductive, autocatalytic citrate cycle (Wächtershäuser, 1992). Hetero-polypeptides with carboxyl- and amide-bearing residues resulted, to form a class of polyanionic, non-replicating biopolymers that broadly incorporated features of the phosphate-based, aperiodic replicators (Westheimer, 1987), consistent with a shared chemistry having shaped the formation of both.

An increase in Asp^1^ and Asn^2^ frequency with phylogenetic depth, on back-tracking to early protein residue sequences, led Brooks et al. (2002) to identify both amino acids as early additions to the code, consistent with their synthesis by 1-to 2-step reaction sequences and inclusion in the first code. Applying changes found to accompany phylogenetic depth to code evolution, within the pre-LUCA era, expanded the scope of standard protein phylogenetics, centered on analysis of post-divergence mutation-distances between residue sequences. In contrast to the foregoing finding, however, homologues, Glu^1^ and Gln^2^, decreased in frequency with phylogenetic depth, suggesting they were late additions to the code. A search for the source of this difference in early levels of incorporation has revealed that Aspartate and Glutamate family amino acids differed in their role in the formation of the code.

Reconciling these path-distance and phylogenetic-depth findings is addressed here with the broad objective of aligning them with the metrics of other traits, to yield a comprehensive reconstruction of code evolution within the pre-LUCA era. They include the dependence of amino acid assignments on codon bonding strength (Lagerkvist, 1978; Grosjean and Westhof, 2016), formation of conspicuous ‘vertical’, mid-base clusters of polar and non-polar amino acids (Woese, 1965; Sonneborn, 1965), codon bias in ancestral sequences (Shepard, 1981; Diaz-Lazcoz et al., 1995; Brooks and Fresco, 2003), homology between cofactor/adaptor tRNA species, with post-LUCA variations removed (Davis, 2008a), and minimization of translation errors. A reconstruction has resulted that depicts the code as a product of the interplay between these factors and that also indicates the source of aminoacyl-tRNA synthetase duality (Eriani et al., 1990).

## 2. Imprint of amino acid synthesis path-distance on the code

Pre-LUCA, tRNA-dependent synthesis pathways (Fig. 1) extending 1- to 2-, 2- to 8-, and 7- to 14-steps, from a precursor in the reductive citrate cycle (RCC), formed amino acids that successively recruited the 5’-, mid-, and 3’-base of encoding RNA triplets. This implies code capacity was 4, 16, and 64 at each stage, leaving, in principle, 60, 48, and 0 triplets unassigned; where unassigned, untranslatable triplets permanently stall translation (Bretscher et al., 1965), in a ‘lethal’ form of nonsense mutation. As the ribosome proofing center allows codons with a 3’U to pair with A, G, C, or U at tRNA N34 (‘wobble’ site), the initial number of untranslatable mutations reduces to 12 mid-base substitutions (XAN: A ➔ C, G, U; X, 5’-coding base; A, mid-site adenosine; N, 3’-any base), making formation of the proposed small, initial code far less hazardous.

**Fig. 1.**
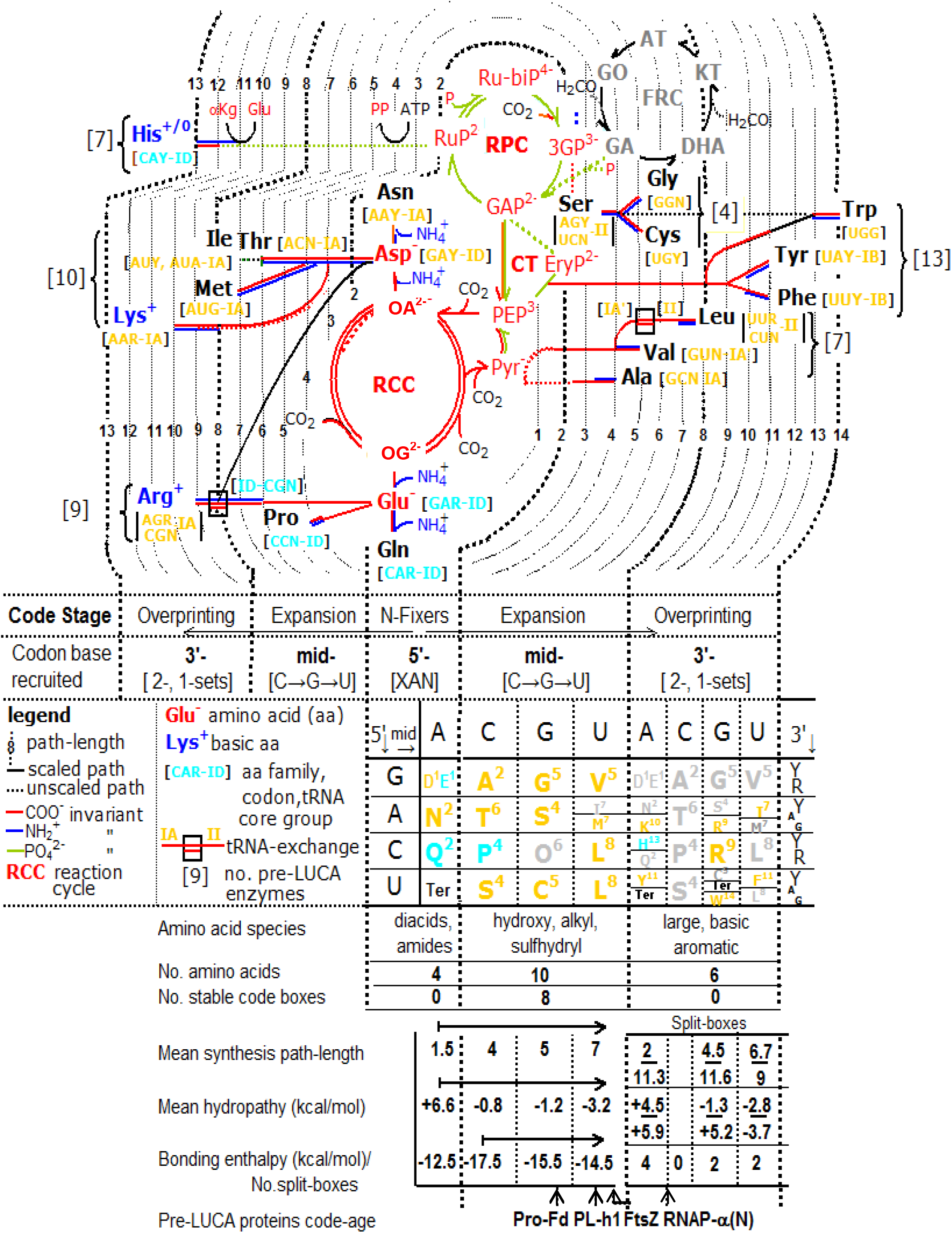
Contour map depicting growth of tRNA-dependent, pre-LUCA amino acid synthesis pathways in relation to code evolution. NH_4_^+^-trapping reactions that branch 1 to 2 steps from 2-oxaloacetate (OA) and oxoglutarate (OG), in the RCC, form Asp^1^ and Glu^1^, and their amides, Asn^2^ and Gln^2^. Other amino acid precursors were from the central trunk (CT) and reductive pentose-phosphate cycle (RPC); FRC refers to the pre-metabolism formose reaction cycle linked to the origin of archaic replicators. NH_4_^+^ fixer amino acids, and Ter signal, acquired codons in the four XAN 4-sets (X, 5’-coding site; N, 3’-degenerate site). Expansion added 10 simple amino acids, formed on 2- to 8-step paths, with increasingly hydrophobic side-chains – hydropathy free energy values are in lower table. Codons were recruited in a compact, mid-C → G → U order, consistent with selection for codon bond strength and error minimization. 3’-Base recruitment added mainly large basic and aromatic amino acids with 9- to 14-step paths. Weak-bonded, pre-assigned boxes were split to form ‘short-path/long-path’ amino acid pairs, and enhanced NAN and NUN hydropathy clusters. Most amino acids occur in the Asp family (gold). Pre-LUCA protein ‘code age’ estimates (below table) are given for pro-ferredoxin, H^+^-ATPase proteolipid helix-1, cell division protein, FtsZ, and RNA polymerase (α-subunit N-terminal). Pro-Fd and PL-h1 ‘code age’ identify surface-contact (Fajan-Paneth effect) and membrane formation as ‘hydrophobic attractors’ in code expansion (Davis, 2002a,b). Reaction sequences (see Supplement, Fig. S1) were drawn from Michal (1999).

Different kinds of amino acids entered the code at each of these three stages: code origin, expansion, and overprinting (Davis, 2007). Diacid amino acids, Asp^1^ and Glu^1^, and their amides, Asn^2^ and Gln^2^, form in NH_4_^+^ trapping reaction sequences that branch 1 and 2 steps from oxaloacetate, or 2-oxoglutarate, respectively. The four NH_4_^+^ fixer amino acids and chain terminator (Ter) acquired triplets in all 4-sets of the XAN column. An invariant mid-A is seen as a vestige of pre-code translation on a poly(A) template. With the participation of an ancestral, type ID tRNA adaptor, charged by either diacid amino acid, and ratchet-equipped ribosome, polyanionic, random sequence poly(Asp^-^,Glu^-^) polypeptides result (Davis, 1999, 2008a,b, 2019).

Expansion from the NH_4_^+^ Fixers Code accompanied recruitment of codon mid-bases: C, G, U. Addition of ten increasingly hydrophobic amino acids, bearing hydroxyl, sulfhydryl, or alkyl side-chains resulted. A columnwise, XCN → XGN → XUN, pattern of codon recruitment added ten amino acids with mean synthesis path lengths of 4, 5, and 7 steps, respectively (Fig. 1). Mid-base recruitment, based on amino acid synthesis path-distances, conforms with selection targeting codon bonding strength (Grosjean and Westhof, 2016) during code expansion.

3’-Codon base recruitment entailed overprinting pre-assigned code boxes. Six long-path amino acids, with basic or aromatic side-chains, were encoded: Arg^9^, Lys^10^, Phe^11^, Tyr^11^, His^13^, and Trp^14^, together with, Ile^7^. Amino acids Lys^10^, His^13^, Trp^14^, and Ile^7^ acquired a codon pair from same family members Asn^2^, Gln^2^, Cys^5^, and Met^7^, respectively. GA^Y^R encodes Asp^1^ and Glu^1^ and as both form on 1-step paths, they depart from the long-path (late-comer)/short-path (early-comer) pattern of code split-boxes, and this is attributed to their joint participation in pre-code translation (Davis, 2019).

Conserved residue profile of pro-ferredoxin (Pro-Fd), H^+^-ATPase helix-1, FtsZ (filamenting temperature-sensitive mutant Z protein), and RNA polymerase (DNA dependent) α-subunit-N-terminus (RNAP-α(N)) were matched with stages identified in code formation, to establish their ‘code age’ within the pre-LUCA era. Pro-ferredoxin is a 23-residue protein (Davis, 2002), reconstructed by Nørgaard (2009) and Nørgaard et al. (2009), with a stage 5.7 residue profile; corresponding to a mid-expansion phase protein. Protein catalyzed RNA synthesis and fluid-mosaic cell membrane proton pumps were seen to be late developments in code expansion (Davis, 2002b).

## 3. Code expansion toward a hydrophobic attractor

Amino acids entering the code at its origin and expansion formed on 1 to 8-step paths that exhibit a length-dependent distribution between code columns: paths of 1.5 ± 0.4, 4.3 ± 0.5, and 6.8 ± 0.6 (m ± s.e.m.) steps occur, respectively, in columns NAN, NCN/NGN, and NUN. These amino acids additionally exhibit a marked path-distance dependent difference in hydropathy. A decrease of −1.5 kcal/mol per step occurs in amino acid side-chain transfer free energy (aqueous to non-polar solvent) with path extension, in the transition, NAN → NCN/GN → NUN (Fig. 2).

**Fig. 2.**
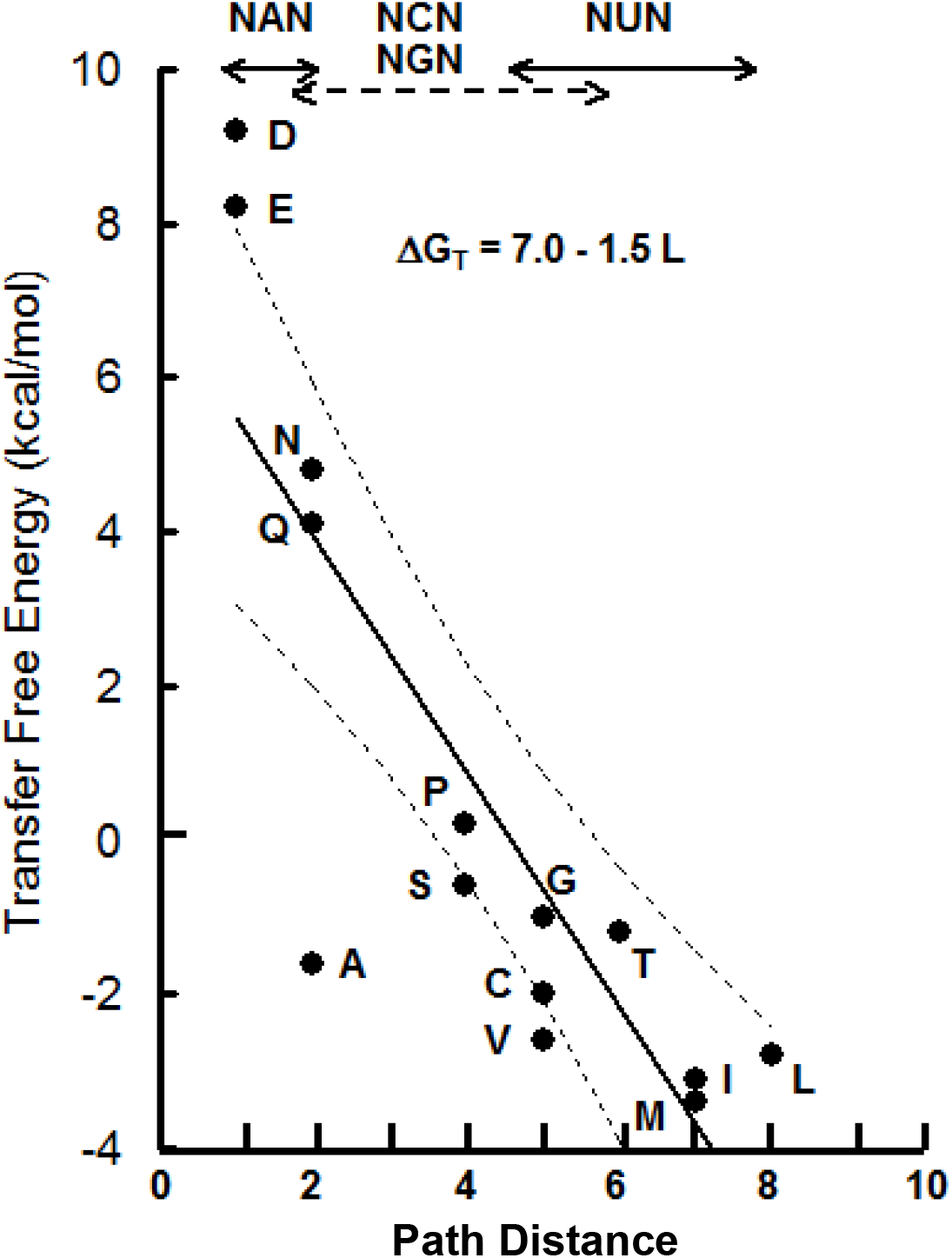
Transition to non-polar amino acids with growth of synthesis pathways during code expansion. Transfer free energy, ΔG_T_ (aqueous to non-polar solvent – dielectric constant 2) values, from Tolstrup et al. (1994), exhibit a linear decrease, −1.5 kcal/mol per step, with increases in synthesis path-distance, L. The 95 per cent confidence interval on the regression line is shown.

From an initial cluster of short-path, hydrophilic amino acids, with NAN set codons, this transition proceeds to a cluster of comparatively long path, hydrophobic amino acids, encoded by NUN set codons. Significantly, it passes through NCN/ NGN encoded amino acids, with transitional hydrophobicity (Supplement, Fig S3), synthesized on paths of intermediate length (Fig. 1). The progressive nature of this transition conforms with the proposed columnwise pattern of expansion (through triplet mid-base recruitment) from a small, first code at NAN. This validates the reliance on amino acid synthetic order, based on path distance, as a measure of advancement in code evolution (Davis, 1999, 2002b, 2011).

It is apparent, from Fig. 2, that the hydropathy clusters in the code, first noted by Woese (1965), are a product, at least in part, of code dynamics during expansion from an initial code with a small set of polar amino acids, encoded by the NAN triplets. The H^+^ ATPase, trans-membrane subunit, proteolipid helix-1 (Davis, 2002b), has a pre-LUCA residue profile matching the amino acid array in a stage-7.1 code (Fig. 1). Synthesis of proteins equipped to partition with a phospholipid bilayer, enabling formation of a fluid-mosaic cell membrane, supplies a hydrophobic attractor for code expansion. An earlier, mid-expansion phase hydrophobic attractor is provided by a low potential ferredoxin antecedent, pro-Fd-[5] (Davis, 2002b, Nørgaard 2009, Nørgaard et al. 2009). Pro-Fd-5 combines a 6-residue (carboxyl-bearing) ‘foot’, <ΔG_T_> = 4.9 ± 2.39 (m ± s.e.m.) kcal/mol per residue, with a 17-residue, [4Fe-4S] cofactor binding segment, <ΔG_T_> = −0.82 ± 0.71 kcal/mol per residue (Supplement, Fig. S2a). Having all anionic residues confined to a ‘foot’, attached to a comparatively long, hydrophobic, cofactor-binding segment, should serve to lengthen attachment to a cationic mineral surface (Fajan-Paneth effect). Pro-Fd[5] thus provides a proto-type for early proteins that functioned as a ‘cofactor anchor’, in a scenario that envisions the genetic code arose within a primordial metabolism, at the surface of a cationic mineral, as envisioned by Wächtershäuser (1992).

## 4. Overprinting of split-boxes by homologous amino acids

Codon G:C and A:U bp frequency at the 5’- and mid-site have long been known to be determinants in the split-box distribution (Lagerkvist, 1978). Figure 3a compares amino acid synthesis path-distances in intact and split-boxes and shows that intact boxes contain short-path amino acids, while, by contrast, split-boxes pair short (early) and long (late) path amino acids. An analysis of variance established that late split-box paths (9.5 ± 1.4 steps) significantly exceed intact, 5.4 ± 0.8 steps, and early split-box paths, 3.6 ± 1.0 steps. This establishes that split-boxes conserve an imprint of when over-printing took place during code evolution. With a short-path amino acid initially assigned an intact codon box, during code origin or expansion, the split-box distribution indicates that, when all 16 code boxes had been allocated, increasing the number of amino acids incorporated into proteins required pre-assigned codons, generally a 3’-purine or 3’-pyrimidine doublet, be reassigned. An incoming long-path (7 to 14 steps) amino acid acquired the codon pair, splitting the box.

**Fig. 3.**
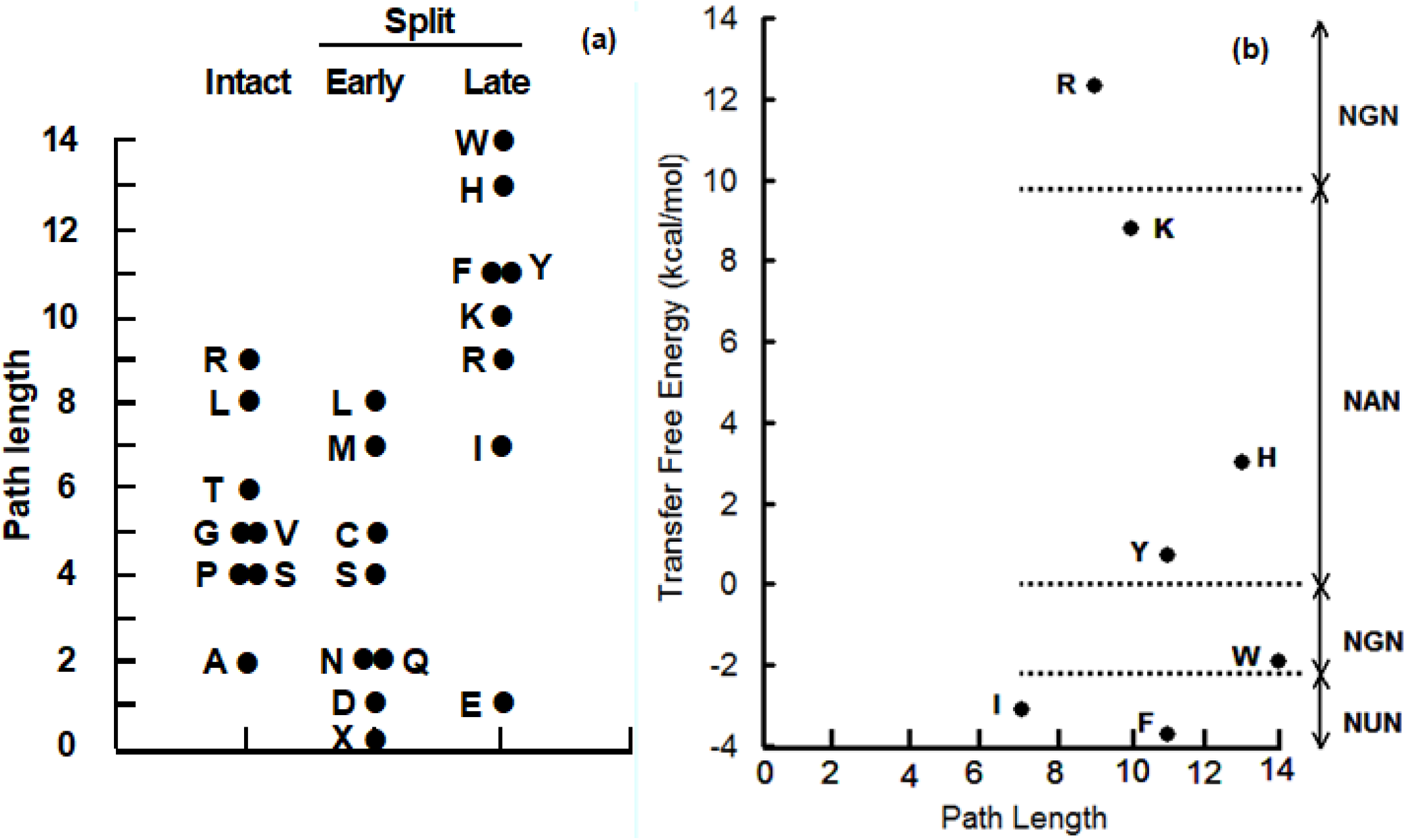
Contribution of path-length dependent over-printing and homology optimization to codon split-box formation. (a) Synthesis path-lengths of intact and split-box amino acids. Early- and Late-set amino acids refer, respectively, the short- and long-path member of each split-box pair. An analysis of variance established path-lengths differed significantly between sets; p = 3.9 x 10^-3^, with means of 5.4 ± 0.8, 3.6 ± 1.0, and 9.5 ± 1.4 (m ± s.e.m.) corresponding to Intact, Early, and Late sets. Late-set paths exceeded both Intact (p < 5 x 10^-2^) and Early paths (< 1 x 10^-3^), in a Tukey’s test; with no significant difference between Intact and Early paths. (b) Shows assignment of homologous amino acids to pre-formed hydropathy clusters, at the codon box-splitting stage (3’-codon base recruitment). An NAN encoded cluster of polar amino acids acquired 3 of 4 polar amino acids, while the NUN cluster added 2 of 3 non-polar amino acids, yielding a probability, p = 5.4 x 10^-3^ of code formation. Single-letter designations include, X, for Ter.

Split-boxes are located within the homology clusters, at NAN (hydrophilic) and NUN (hydrophobic) code columns, to a statistically significant degree (Fig. 3b). All split-box codon sets (AA^Y^_R_, GA^Y^_R_, CA^Y^_R_, UA^Y^_R_) (AU^YA^_G_, UU^Y^_R_), within the hydropathy clusters were distributed to homologous amino acids, (Lys^10^/Asn^2^, Asp^1^/Glu^1^, Tyr^11^/Ter, Gln^2^/His^13^) (Ile^7^/Met^7^, Phe^11^/Leu^8^), discounting (UA^Y^_R_, Tyr^11^/Ter). From this perspective, the code is optimal, with a joint distribution probability of 5.4 x 10^-3^.

Acquisition of AGR and UGG, by Arg^9^ and Trp^14^, respectively, resulted in non-cluster split-boxes. Split boxes also occur within code domains – containing amino acids with the same precursor, related tRNA species, and contiguous codons (Davis, 2008a,b). A factor directly linked to Lagerkvist’s rule was plainly at play in forming the split-box distribution. When reassignment of a codon doublet was favorable, at an advanced stage in code formation, amino acid homology was optimized, consistent with minimizing the deleterious effects of amino acid substitution, as Sonneborn (1965) envisioned.

## 5. Code enthalpy states and the split-box distribution

Codon bonding strength played a key role in code formation, as illustrated by its split-box distribution (Lagerkvist, 1978). Free energy determinations from the thermal dissociation of paired nucleotide dimers (Chen et al., 2012) were seen by Grosjean and Westhof (2016) to furnish a physical parameter linked to translation fidelity and, plausibly, to the order of codon recruitment during code evolution. Figure 4a depicts the distribution of three possible H bonds, in weak, mixed, and strong bp dimers. Enthalpy values, from the expanded set of Chen and coworkers (see Supplement, Table S2) decrease directly with the number of dimer N-H••O bonds, at a rate of −2.73 kcal/mol per bond. Based on mid-range values in H bond interaction energy distributions obtained by Wendler et al. (2010), the comparative N-HooN and and C-H••O bond enthalpy, in an isobaric, aqueous medium, is found to be −1.72 and −0.96 kcal/mol, respectively (Table 1).

**Fig. 4.**
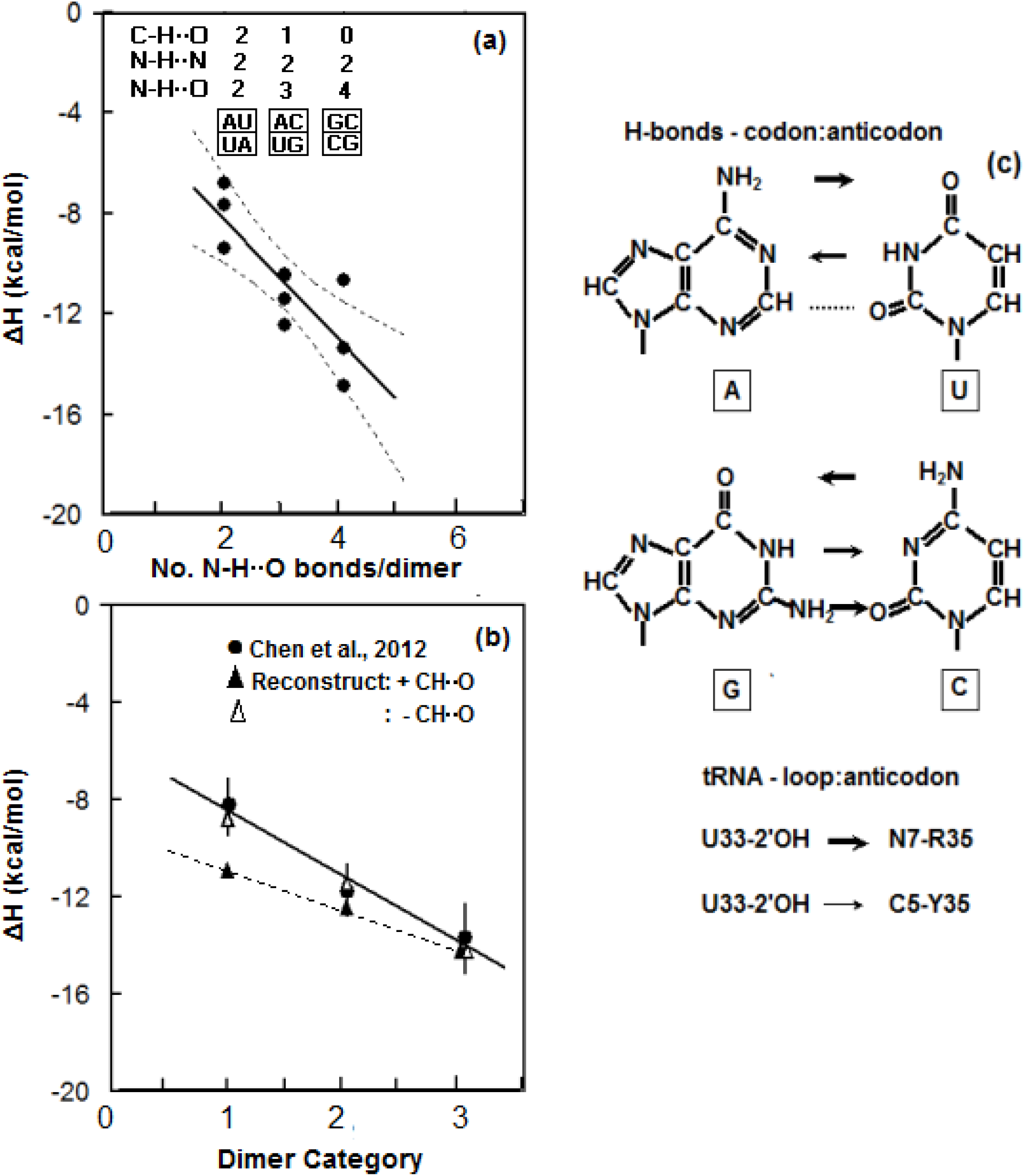
Codon H bond strength in translation. (a) Bond formation enthalpy, from thermal stability of complementary nucleotide dimers (Chen et al., 2012; expanded set), is shown to decrease linearly with the number of N-H••O bonds, n: ΔH = −3.09 – 2.73n, depicted within a 95 per cent confidence interval and goodness of fit. (b) Estimated N-H••O, N-H••N, and C-H••O bond enthalpy (Table 1), in aqueous phase, reveal dimers with weak, A:U, mixed A:U, G:C, or strong, G:C, bp have aggregate N-H••O and N-H••N enthalpy of −8.2, −11.6, and −14.4 kcal/mol, respectively. These values are within the observed standard error for each category mean: weak, −8.2 ± 0.60; mixed, −11.8 ± 0.53; and strong, −13.7 ± 1.51 kcal/mol (refer to Supplement, Table S1). In a conspicuous departure from observed enthalpies, inclusion of C-H••O enthalpy produced values of −11.0, −12.6, and −14.4 kcal/mol per dimer with weak, mixed, and strong bonds, respectively. (c) Depicts codon 5’- and midbase H-bonds within the A-type double-helix formed with anticodon bases B36 and B35. tRNA loop nucleotide, U33, additionally forms an N-H••O bond with anticodon purine, R35, or C-H••O bond with pyrimidine, Y35. Arrows indicate H bond direction and strength: ➞ 2.73, ➞ – 1.72, ➞ −0.96 kcal/mol.

**Table 1.** Estimated H bond enthalpy

| H Bond | Interaction Energy<sup>1</sup><br>(kcal/mol) | Ratio | Estimated Enthalpy<sup>2</sup><br>(kcal/mol) |
| --- | --- | --- | --- |
| N-H••O | $-2.48 \pm 0.90$ | 1.0 | -2.73 |
| N-H••N | $-1.55 \pm 0.33$ | 0.63 | -1.72 |
| C-H••O | $-0.87 \pm 0.38$ | 0.35 | -0.96 |

<sup>1</sup> Mid-range H bond interaction energy values, within half-range limits, from Wendler et al. (2010).
<sup>2</sup> N-H••N and C-H••O enthalpies estimated from energy ratios applied to N-H••O enthalpy, using the dependence of nucleotide dimer-pair formation enthalpy on N-H••O depicted in Fig. 4(a).

Figure 4b shows the enthalpy for weak, mixed, and strong dimers, obtained by combining N bearing H bond estimates, from Table 1. They closely agree with those reported by Chen and coworkers. Inclusion of C-H••O enthalpy results in a conspicuous departure from observed dimer bp bonding values, particularly in weak dimers. Thermodynamic evidence thus excludes a third H bond at [2C-H••O=C2], on the A:U Watson-Crick face (Fig. 4c): where double-helix bp distances are 2.8 Å for N-H••O, 3.0 Å for N-H••N (Klug, 2004), and 3.2 Å at [C-H••O]. As Fig. 4c indicates, a 2’O-H••C5 bond forms between an invariant tRNA anticodon loop U33 and anticodon Y35. Alternatively, a strong 2’O-H••N7 bond is formed with a purine at N35 (Grosjean and Westhof, 2016).

Lagerkvist’s split-box ‘two out of three’ rule can be restated in terms of codon box enthalpy states. Symmetry between the 5’-site of strong, G/C v. C/G, and weak, A/U v. U/A, bonders yields equal enthalpy, reducing the number of distinguishable codon box states by half (Fig. 5a). A second symmetry arises with compensating 5’- and mid-site differences in mixed-strength codon boxes, as [G/C, A] ≡ [A/U, G], and [G/C, U] ≡ [A/U, C], decreasing the number of distinct code box states to six:

|  |  |
| --- | --- |
| (1) 5S + 2M = -19 kcal/mol | [G/C],C; Ala<sup>2</sup>, Pro<sup>4</sup> |
| (2) 4S + 2M + W = -17 | [G/A],G; Gly<sup>5</sup>, Arg<sup>9</sup> |
| (3) 4S + 2M = -16 | [G/C],U; Val<sup>5</sup>, Leu<sup>8</sup><br>[A/U],C; Ser<sup>4</sup>, Thr<sup>6</sup> |
| (4) 3S + 2M + W = -14 | [G/C],A; Asp<sup>1</sup>/Glu<sup>1</sup><br>Gln<sup>2</sup>/His<sup>13</sup><br>[A/U],G; Ser<sup>4</sup>, Arg<sup>9</sup><br>Ter/Cys<sup>5</sup>/Trp<sup>14</sup> |
| (5) 3S + 2M = -13 | [A/U],U; Met<sup>7</sup>/Ile<sup>7</sup>, Leu<sup>8</sup>/Phe<sup>11</sup> |
| (6) 2S + 2M + W = -11 | [A/U],A; Ter/Tyr<sup>11</sup><br>Asn<sup>2</sup>/Lys<sup>10</sup> |

where S = −3, M = −2, W = −1 kcal/mol; whole number equivalents of H bond enthalpy in Table 1.

**Fig. 5.**
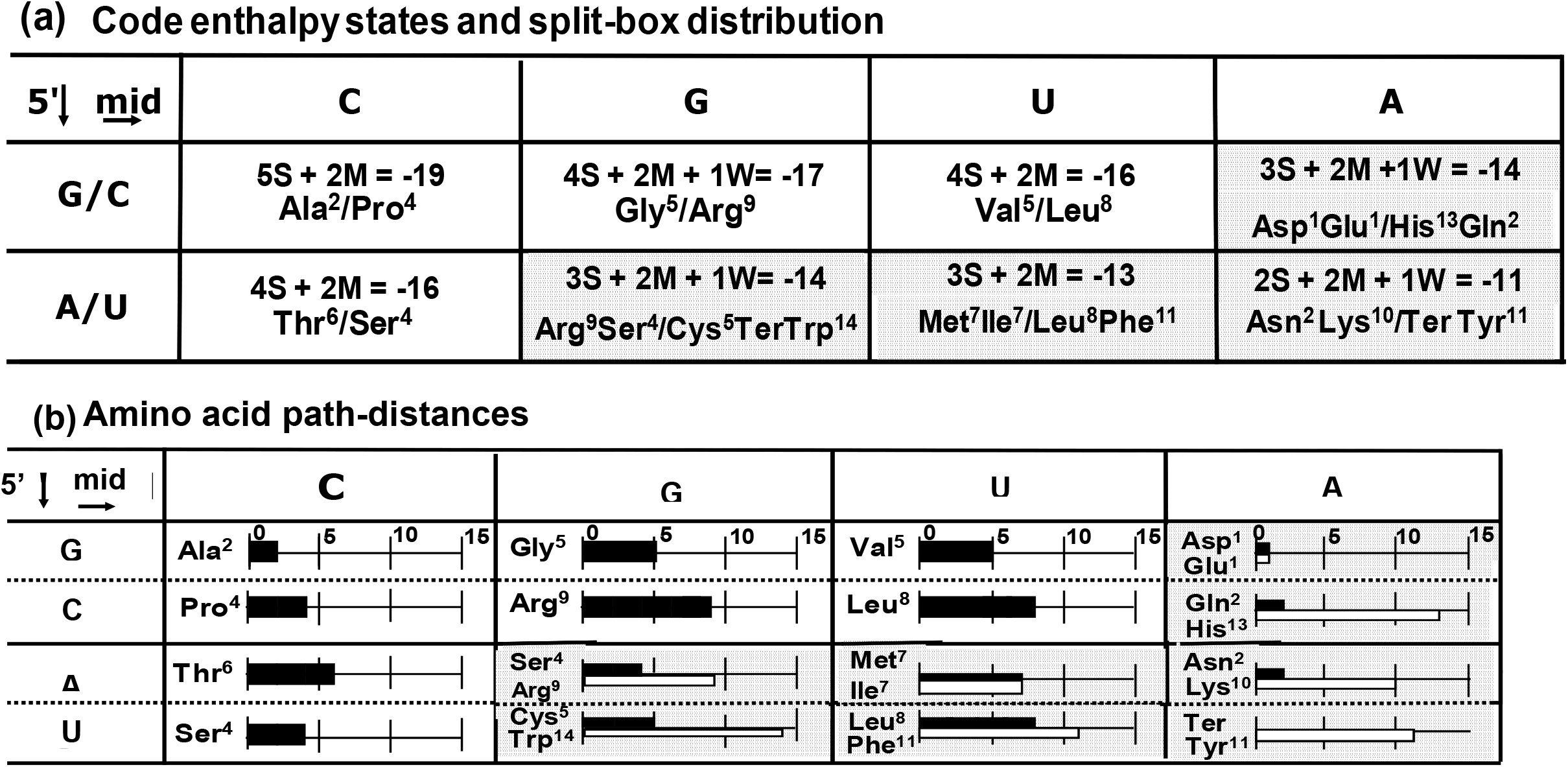
Codon split-box distribution in relation to proofing center enthalpy states and amino acid synthesis path-length. (a) G ↔ C and A ↔ U equivalence reduces the number of distinct, 5’-site H bond enthalpy states to two. tRNA anticodon loop U33 introduces a mid-site asymmetry, resulting from a strong 2’O-H••N7 bond with anticodon purine, R35, and weak 2’O-H••C5 bond with pyrimidine, Y35. Four distinguishable mid-site states result. The eight doublet states obtained are defined by six equations, where variables S, M, and W denote strong, moderate, and weak H bonds; respectively, N-H••O, NH••N, and C-H••O. With rounded values of −3, −2, and −1 kcal/mol (Table 1), all four weak codon states, bonding enthalpy exceeding −16 kcal/mol, are split (stippled boxes). (b) All intact boxes have short-path (up to 8 steps) amino acids (superscripts/bar show path-length), 6 of 8 split-boxes encode amino acids with paths of 7 to 14 steps (white bar), making box splitting a late event.

States with codon:anticodon proofing center bonding enthalpy higher than −16 kcal/mol are split in the standard code. Split boxes pair amino acids with synthesis paths of mixed length (Fig. 5b), as noted in Sec. 3. Codon boxes with sub-threshold bonding strength were, consequently, overprinted late in code formation.

## 6. Compact transitional codes minimized risk of untranslatable triplets

Figure 6 illustrates a pattern-dependent difference in the risk of mutation to an unassigned, untranslatable template triplet, during formation of the code. Until a triplet acquires an adaptor, it is unreadable. Mutation to an unassigned triplet in a transitional code can, consequently, permanently stall translation (Bretscher et al., 1965). Minimizing the risk of this ‘lethal’ form of nonsense mutation could be expected, therefore, to have strongly shaped the initial pattern of codon assignments, when the number of encoded amino acids and their cognate cofactor/adaptor tRNA was still small. Thus, the compact ‘four amino acid’ NH_4_^+^ Fixers Code, is 25 per cent less susceptible to translation-halting nonsense mutations than the semi-compact code code of four amino acids maximizing codon bonding strength, within the ribosome proofing center (Grosjean and Westhof, 2016); the mean enthalpy difference between both codes being −5.5 kcal/mol (−12.5 vs.-18 kcal/ mol). Adaptor:template bonding strength underlies template decoding specificity. It is not surprising, therefore, that path-distances indicate expansion from the NH_4_^+^ Fixers Code resulted in amino acids, with a mean synthesis path-length of 4 steps, saturating the NCN column triplets, enabling a near-maximal mean bonding enthalpy of 17.5 kcal/mol. In addition, by retaining a compact pattern of codon assignments, a one-third reduction in translation-halting, ‘Bretscher’ mutations resulted.

**Fig. 6.**
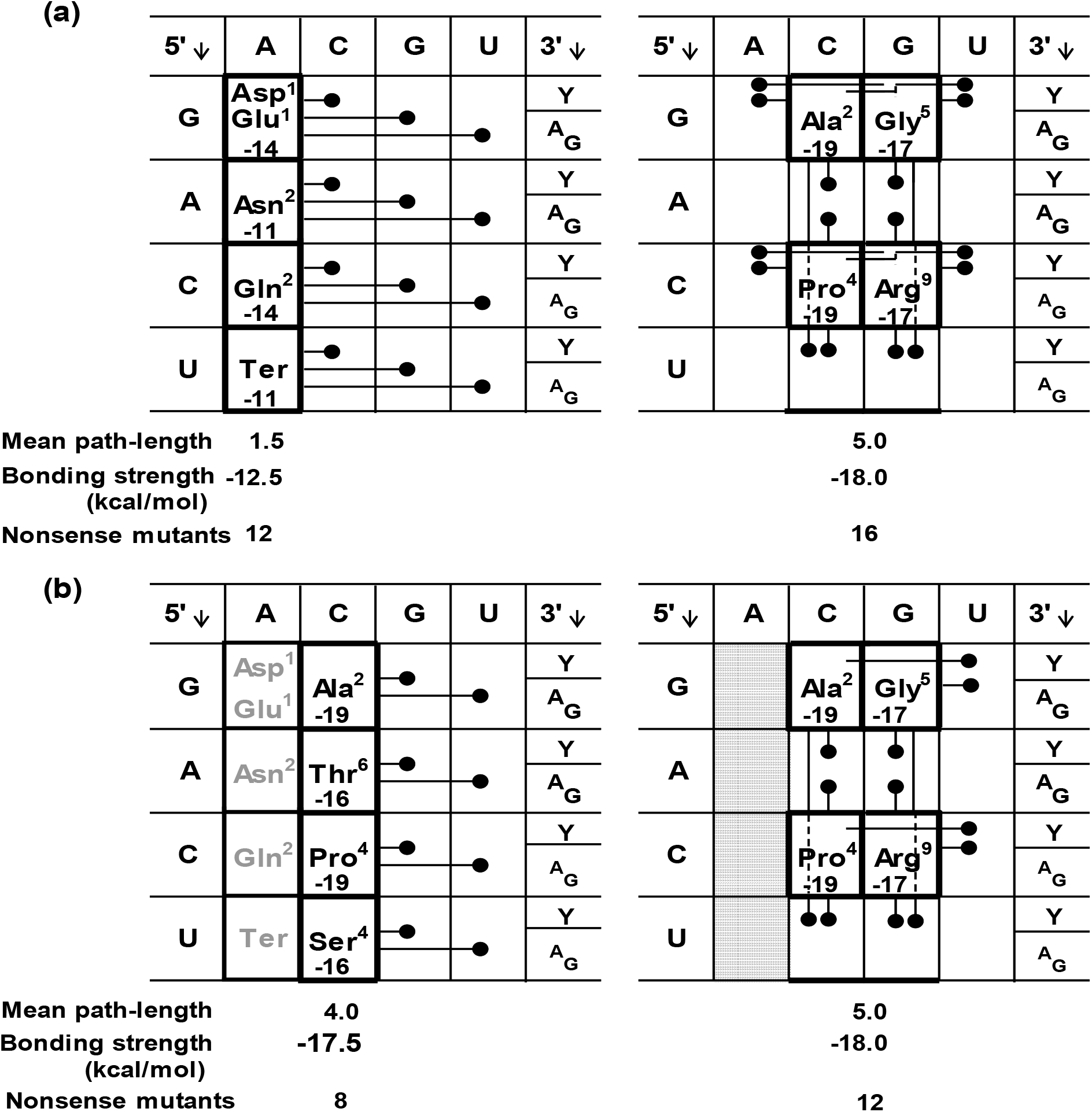
Minimizing amino acid synthetic order offered a preferred track for code evolution. (a) NH_4_^+^ Fixers Code with four, 1- to 2-step synthesis path amino acids and a Ter signal, saturated the NAN codon set. This lowered by one-quarter, from 16 to 12, the number of possible nonsense mutations, to an unreadable, translation-halting triplet, in comparison with encoding the four amino acids optimizing mean codon bond enthalpy, at −18 kcal/mol, with a semi-compact distribution of 2- to 9-step amino acids. (b) Expansion of the NH_4_^+^ Fixers Code to an equally compact NCN codon set with four, 2- to 6-step (8 amino acids, total) amino acids achieves near-optimal codon bonding, at −17.5 kcal/mol, with a one-third, from 12 to 8, lower rate of translation-halting mutations; indicated as, 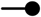.

With mean synthesis path-lengths extending 4, 5, and 7 steps per codon set, expansion beyond the NH_4_^+^- Fixers Code, at NAN set, proceeded in an NCN → NGN → NUN order. Mean codon bond enthalpy accompanying expansion was −17.5, −16.1, and −14.5 kcal/mol per set (Fig. 1). This provides evidence substantiating that selection optimized translation fidelity, through assigning codons of highest bonding strength first (Grosjean and Westhof, 2016). With the threat posed by unassigned triplets greatest initially, and receding as code expansion advanced, assignment of weak-bonding NAN codons, with mean bonding enthalpy of −12.5 kcal/mol (Fig. 6) is understandable, given that amino acid synthetic order, based on path-distances, reveals the first code arose from pre-code translation on a poly(A) template, precluding competition with NCN, NGN and NUN codon sets.

Columnwise organization of the code (Fig. 1), first noted by Woese (1965), is seen here as the outcome of an interplay between its origin, from pre-code assembly of a few amino acids on a poly(A) template, the risk posed by untranslatable codons, and selection for homology and codon bonding strength.

## 7. Imprint of pre-code translation on the code

The two diacid amino acids in proteins share the GA^Y^_R_ box, forming the only short-path pair to occupy a split-box. With a mean bond enthalpy of −14 kcal/mol (Fig. 5a), GA• box codons would split during 3’-base recruitment, in the final stage of code formation (Sec. 2). Each forms on a 1-step path, and as first generation additions to the code, Asp^1^ and Glu^1^ conceivably shared a cofactor/adaptor, tRNA-!D^Asp,Glu^_3’CUU_, cognate for GA• codons, in the manner of pre-code translation of both amino acids on a poly(A) template (Davis, 1999, 2019). Formation of the NH_4_^+^ Fixers Code, with the acquisition of codons in the XAN set, and subsequent expansion to XCN set codons are reconstructed in Fig. 7, from amino acid path-distances and homology between pre-divergence tRNA base sequences (Davis, 2008a,b).

**Fig. 7.**
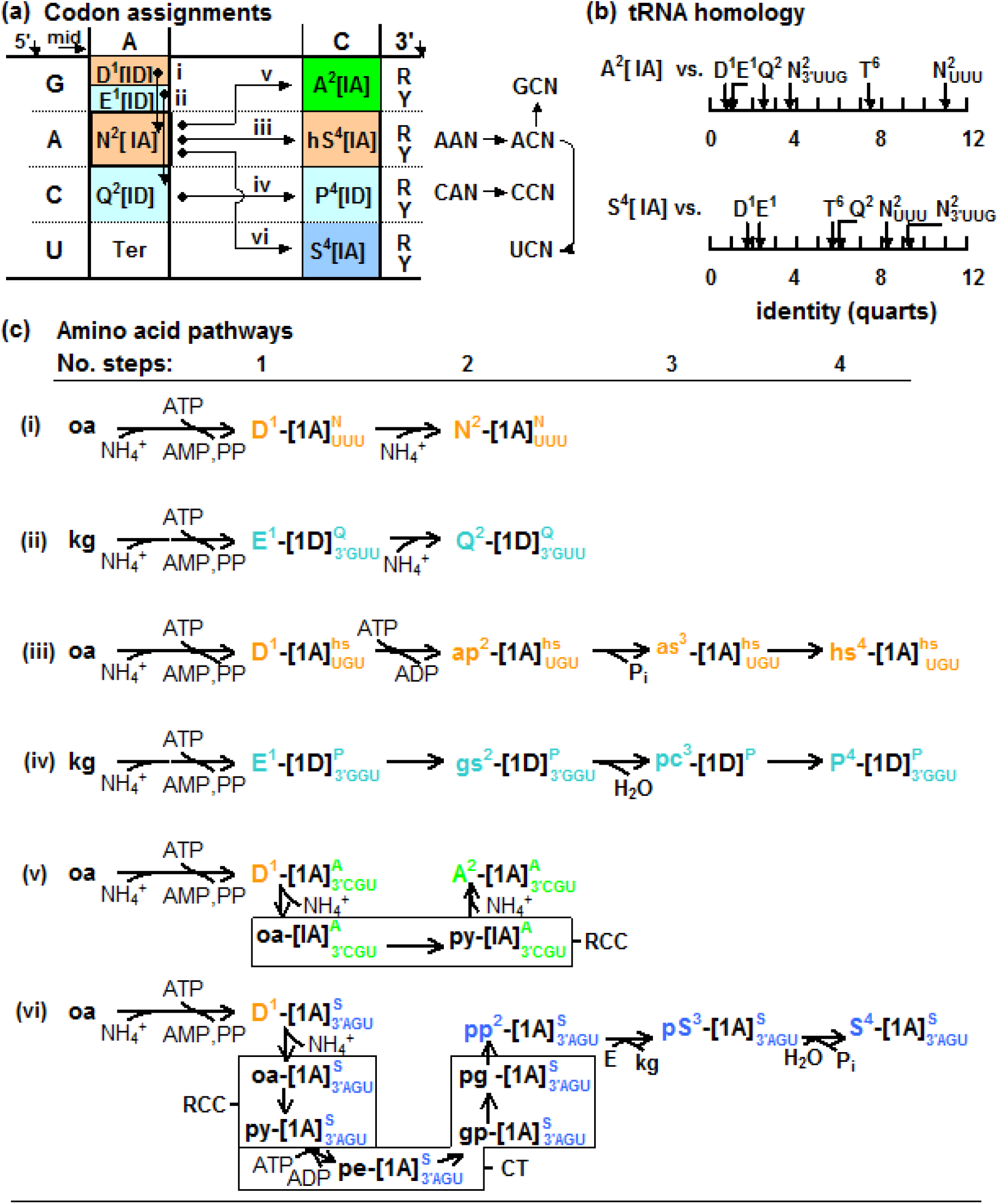
Pathway extensions in expansion from the NH_4_^+^ Fixers Code. They accompanied codon acquisition by newly formed amino acids and diversification of LUCA tRNA species. (a) Initial NH_4_^+^ fixing reactions (paths i and ii) formed diacid amino acids, D^1^, E^1^, and their amides, N^2^, Q^2^. Together with Ter triplets, they are shown to have recruited all 16 NAN codons. With 5’-base identity determining codon specificity in the NH_4_^+^ Fixers Code and 3’-base degeneracy, mutation to any of 48 unassigned triplets, able to halt translation, decreases from ‘3 in 4’ (random array) to ‘1 in 3’ (mid-A substitution). Recruiting an NCN template triplet required a mid-base, A➔ C (adaptor U35 ➔ G35) substitution, Extension of amide synthesis pathways, iii and iv, yield 4-step paths producing a Thr^6^ α-amino acid intermediate, homo-serine (hs^4^), and imino acid, proline (P^4^), respectively. Subsequent 5’-base template substitutions, A ➔ G and A ➔ U, in hS^4^ codons, ACN, form codons GCN and UCN for A^2^ and S^4^, synthesized on paths-v and -vi, respectively, saturating the 16 NCN set. All transitions conserve cofactor/adaptor tRNA core structure (in brackets); apart from D^1^[ID] ➔ N^2^[IA]. (b) Consensus sequences of LUCA tRNA specific for A^2^ and S^4^ exhibit homology of 7.4 to 11.0 quarts (p = 3.5 x 10^-5^ – 2.4 × 10’^7^) and 5.7 to 9.2 quarts (p = 3,7×10^-4^ – 2.9×10^-6^) with tRNA for N^2^ and T^6^ (T^6^ being terminal amino acid of present hS^4^ path). (c) Synthesis path-i to -iv have linear extensions. Path-v and -vi result in A^2^ and S^4^ synthesis and they include takeover of pre-code, supernumerary reaction sequences from RCC and CT, together with N^2^ pathway extensions. Letter and background colors indicate synthesis pathway precursor – oa, oxaloacetate; og, oxoglutarate; branch-point - py, pyruvate; pg, 3-phospho-glycerate.

Initial saturation of the sixteen XAN triplets (Davis, 1999), recruited the weakest set in the code (mean codon bonding enthalpy, −12.5 kcal/mol; Fig. 5). Formation of the first code plainly predated competition with strong bonding codons and translation at optimal fidelity. With translation originating on a simple, pre-code poly(A) template, resulting in synthesis of polyanionic, random sequence poly(Asp, Glu) polypeptides, the transition to a set of codons with an invariant mid-A indicates formation of the NH_4_^+^ Fixers Code represented an intermediate stage in the transition to a code; insufficiently advanced to have optimized codon bonding strength. Expansion from the weak NAN triplets of the NH_4_^+^ Fixers Code, by contrast, displays selection for codon bonding strength, in full force. As shown in Fig. 7, an equally compact code, incorporating NCN triplets – the strongest bonding codon column (bond enthalpy, −16 to −19 kcal/mol; Fig. 5) resulted.

NH_4_^+^ Fixers Code amino acids derive from oxoglutarate (Glu^1^, Gln^2^), or oxaloacetate (Asp^1^, Asn^2^). While Asn^2^ has a type-IA tRNA, Asp^1^, Glu^1^, and Gln^2^ have type-ID tRNA (Saks and Sampson, 1995). Hence, a type ID tRNA core was the apparent ancestral form (Davis 2006, 2008). With its A-rich codons, the NH_4_^+^ Fixers Code indicates pre-code translation occurred on a poly(A) template, making AAA, the ancestral codon. Since AAA codes for Asn^2^, in the NH_4_^+^ Fixer Code (assignment of AAA to Lys^10^ in the standard code followed from a later overprinting), the distinct core structure of tRNA-IA_Asn__UUU_ can be linked, accordingly, to direct competition for the AAA codon and elimination of the ancestral, ambivalent tRNA-ID^Asp,Glu^_AAA_ cofactor/adaptor; together with replacement of codon:anticodon self-recognition by base pair complementarity (Davis, 2019).

Unlike the linear i – iv paths (Fig. 7c) yielding Asn^2^, Gln^2^, homo-Ser^4^, and Pro^4^, respectively, paths v and vi show oxaloacetate as an upstream precursor in pathways leading to Ala^2^ and Ser^4^ synthesis (Davis 2013). Close homology between pre-divergence tRNA-IA^Ala^_3’CGU_ and tRNA-IA^Ser^_3’CGU_ (pre-type II form) versus tRNA-IA^Asn^_UUU_ (Fig. 7b) suggests oxaloacetate ➔ pyruvate, and oxaloacetate ➔ phosphoglycerate reaction segments (with carboxyl-bearing intermediates for cofactor tRNA attachment) were deleted during the protein takeover of amino acid synthesis. As segments upstream from current path branch-points include central metabolism reactions, predating amino acid synthesis, they were supernumerary to the path-distance metric of code evolution; incorporation of Val^5^ synthesis reactions in the Ile^7^ pathway (Michal, 1999) illustrates a path takeover in code formation. Homology between pre-species-divergence tRNA species for Ser^4^ and Asn^2^, indicate type-II tRNA^Ser^, bearing an enlarged variable loop, evolved some time after tRNA-IA’^Cys^_3’ACU_ had formed from its precursor cofactor/adaptor, tRNA-IA^Ser^_3’AGU_.

Emergence of tRNA-II^Ser^ allowed transfer of anticodon stem identity elements to the variable loop, freeing the adaptor to diversify the number of Ser^4^ codons, at a time when unassigned, untranslatable triplets posed an existential threat to protein synthesis. Codons initially recruited for Ser^4^, by variant tRNA^Ser^ species, were evidently acquired by Cys^5^, Gly^5^, Leu^7^, Arg^9^, Trp^14^, and Ter. In particular, α-isopropyl-malate, a dicarboxyl intermediate in Leu^7^ synthesis, is the putative site of a tRNA exchange, resulting in Leu^7^ acquiring a type-II tRNA species from Ser^4^ – its sole source – cognate with UUN and CUN (Davis, 2013).

tRNA sequence homology, preceding species divergence (Davis, 2008a,b), reveals three of the four amino acids saturating the set of strong NCN codons, on expansion from the NH_4_^+^ Fixers Code, had type-IA cofactor/adaptors (Fig. 1a,b). They include the 4-step α-amino acid, homoserine, hS^4^, an intermediate in the synthesis of Thr^6^; end-product of the full, 6-step path and standard set amino acid. Single N36 anticodon U ➔ C and U ➔ A substitutions in tRNA-IA^hS^_3’UGU_, respectively, recruited codons GCN and UCN for Ala^2^ and Ser^4^. Anticodon N35 U ➔ G substitutions in tRNA-IA^Asn^_UUU_ and tRNA-ID^Gln^_3’GUU_, respectively, occurred in forming tRNA-IA^hS^_UGU_ cognate with codons ACN, and tRNA-ID^Pro^_3’GGU_ cognate with codons CCN. In expansion from the NH_4_^+^ Fixers Code, codon bonding strength increased substantially, as indicated, from −12.5 to −17.5 kcal/mol (Fig. 5a), while mean amino acid synthesis path-length increased from 1.5 to 3.5 steps, and mean accumulated codon substitutions rose from 0.5 to 1.5. The IA:ID ratio in cofactor/adaptors with each core structure changed from 1:2 (Asp^1^ and Glu^1^ ambiguously charging tRNA-ID^Asp,Glu^_3’CUU_) to 3:1. Since this measure broadly indicates the relative proportion of amino acids in the Asp and Glu family within the code, it demonstrates code expansion resulted from disproportionate growth of the Aspartate family.

## 8. Correlation between amino acid path-distance and phylogenetic depth

Determinants of the order of entry of different kinds of amino acids into the code, and their distribution within it, have been identified above using the synthetic order of amino acids, based on path-length, as a metric of the advancement of code evolution. An alternative measure of the course of code evolution has been introduced that relies on amino acid phylogenetic-depth (Brooks et al., 2002). It extends standard phylogenetic techniques of protein analysis to sort amino acids into early-, mid-, or late-additions to the code, based on the direction and magnitude of change in residue frequency, on back-tracking to predivergence protein residue sequences In principle, both metrics of code evolution would be anticipated to align.

Figure 8a shows there is a linear correlation between path-length and phylogenetic-depth among standard set amino acids, on excluding the four Glutamate family members: L = 13.7 – 7.5 D, where L denotes pathlength and D is depth. With depth specified as the ratio of probabilities, p_LUCA_(i)/p_mod_(i), for residue, i, in a LUCA era protein versus the modern protein, code earlycomers Asp^1^, Asn^2^, and Ala^2^ display, for example, depth values of 1.06 to 1.43, while latecomers Phe^11^, Tyr^11^, Trp^14^ depth is 0.59 to 0.35 (Supplement Table S2). Differences in amino acid path-length, within the three depth categories of code formation (1, early; 2, mid; 3, late) were statistically significant, with probability, p = 3.2 x 10^-3^ (F = 8.93, d.f. = 2), in an analysis of variance; early and late amino acid mean path-distances differed significantly in a Tukey’s test, p < 1 x 10^-2^.

**Fig. 8.**
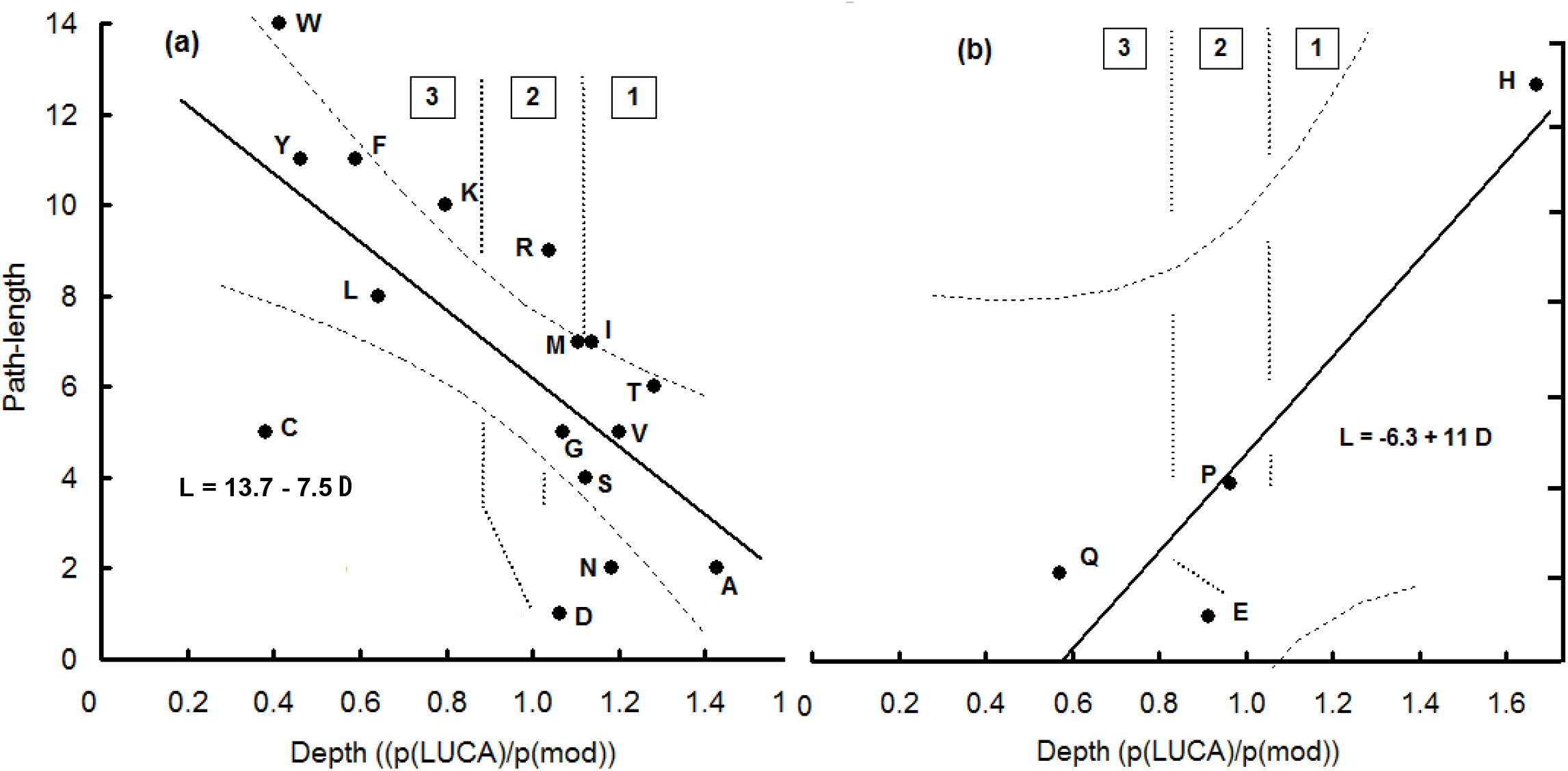
Linear regression, with 95 per cent confidence interval, showing a contrasting relationship between synthesis path-length and phylogenetic-depth in each amino acid family. (a) Asp family, n = 16, and (b) Glu family, n = 4. Regions 1, 2, and 3 refer, respectively, to higher (early-comer), constant (mid-stage), and lower (late-comer) depth index for indicated residues in LUCA versus modern proteins (Supplement, Table S2). L signifies synthesis path-length and D phylogenetic-depth in the regression equations.

Phylogenetic depth and amino acid synthetic order were, conversely, negatively correlated in the Glutamate family, containing Glu^1^, Gln^2^, Pro^4^, and His^13^: L = −6.3 + 11.0 D. This incongruously places both short-path NH_4_^+^ fixer amino acids, half the NH_4_^+^ Fixers Code, among late-comers (category 3) to the code, while Pro^4^ and His^13^ were, respectively, categorized as intermediate (category 2), and early (category 1) additions to the code. The Glutamate Family is comparatively small, only one-fifth of the standard set, and this will be seen to be significant, together with phylogenetic evidence, noted in the previous section, on the diversification of bifunctional cofactor/adaptor tRNA species in the pre-LUCA era.

The quantitative agreement demonstrated between path-distance and phylogenetic-depth validates both as metrics of code evolution. It remains to be clarified, however, why they should be aligned for amino acids with oxaloacetate, pyruvate, phosphoglycerate, and phosphoenolpyruvate as precursors, but misaligned for those derived from oxoglutarate.

## 9. Antiquity of NH_4_^+^ fixer amino acids and their tRNA

Prokaryotes conserve tRNA-dependent pathways for the synthesis of both diacid/amide pairs of NH_4_^+^ fixer amino acids (Wilcox and Nirenberg, 1968; Schon et al., 1988; Curnow et al., 1996; Gagnon et al., 1996). This supports them being the conserved root of a once extensive network (Cork and Purugganan, 2004) of tRNA-dependent amino acid synthesis pathways. Both NH_4_^+^ fixer diacid/amide pairs thus formed before most standard set amino acids.

Figure 9 depicts a tree of tRNA species based on homology between consensus base sequences, predating species divergence (Davis, 2008a,b). Its relation to code evolution is inferred from the path-distance synthetic order of each cognate amino acid. All four NH_4_^+^ fixer cofactor/adaptor tRNA species cluster at the tree root, consistent with forming the first code. Other amino acids, with longer, later, and less-integrated synthesis pathways, evidently discarded their tRNA cofactor during the protein takeover, with tolerable disturbance to the network of synthesis pathways.

**Fig. 9.**
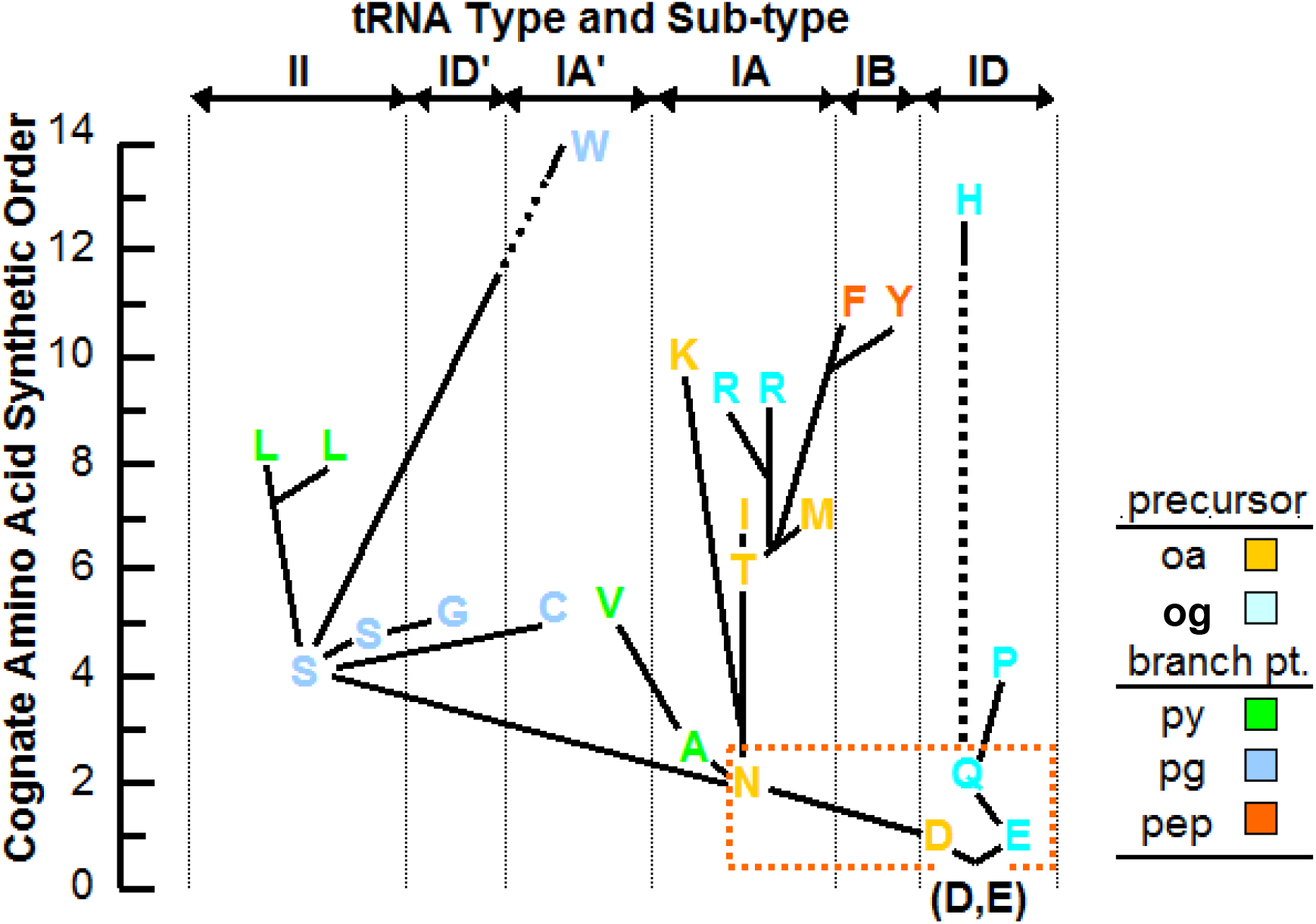
Phylogenetic tree of LUCA tRNA species showing substantially greater diversification of Asp family tRNA, from a root of diacid and amide amino acid adaptor/cofactors. Nearest-neighbor tRNA having closest identity between consensus sequences, excluding post-LUCA variations and universally conserved nucleotides, form each node. Cognate amino acid synthesis path-length, from the reductive citrate cycle, specifies the stage of code formation at each node, marking the origin of a tRNA species. Fourteen amino acids, encoded by 17 sets of codons, charge tRNA derived from Asn^2^ tRNA. Two amino acids, each with a single set of codons, charge tRNA derived from tRNA^Gln^. Enclosed at the tree root are tRNA for both diacid amino acids (D, E) and their amides (N, Q), highlighting their retention of tRNA-dependent amino acid synthesis in prokaryotes. This supports their being the conserved root of an early network of tRNA-dependent amino acid synthesis pathways. Each pathway precursor, or branch point, in central metabolism is indicated. A discontinuous line indicates intermediates lack an α-carboxyl for cofactor/adaptor tRNA attachment. Leu^8^ and Arg^9^ pathways contain a dicarboxyl intermediate, providing a site for tRNA exchange, resulting in anomalous adaptor and codon assignments. An over-bar shows type/sub-type of tRNA for each amino acid, using Saks and Sampson (1995) classification. Letter colors mark amino acid precursor – oa, oxaloacetate; og, oxoglutarate; and branch-point – py, pyruvate; pg, 3-phospho-glycerate; pep, phosphoenolpyruvate.

Tree nodes join each tRNA species and its antecedent, identified from base sequence homology. Cognate amino acid path-length determined branch height (Fig. 9). Conspicuously more, 14 versus 2, tRNA species diversified from tRNA-IA^Asn^ than tRNA-ID^Gln^. Although Glu^1^ is precursor to Gln^2^, Pro^4^, and Arg^9^, a tRNA exchange at a triple carboxylated intermediate, arginino-succinate (step-8), is credited with Arg^9^ acquiring an Asn family cofactor/adaptor, with a type-IA core structure, yielding isoacceptors cognate with codons AGR and CGN in Asn^2^ and Gln^2^ code domains, respectively (Davis, 2013). With His^13^ synthesis originating at D-ribose-5-phosphate, the amino acid did not plausibly acquire a Glu cofactor/adaptor until histidinol-phosphate synthesis, 2 steps prior to His^13^ formation.

Thus Gln^2^ and Pro^4^ are the only fully tRNA-dependent members of the Glutamate family. It is an open question, therefore, whether the 13 step His synthesis path provides a reasonable measure of its time-of-entry to the code. As a first generation addition to the code, the CAN box initially coded for Gln^2^ in the NH_4_^+^ Fixers Code. Acquisition of the CAY doublet by His^13^ places its addition to the code in the final stage of its formation (Fig. 1). Early His^13^ entry to the code could not, in view of this, be the source of its 1.67 depth index, as reported by Brooks et al. (2002). Although it is apparent, from Fig. 8a, that depth measurements convey information on amino acid time-of-entry into the code, Glutamate family results suggest it is responsive to other factors influencing pre- and post-LUCA protein residue frequencies.

Figure 10 depicts the difference in cumulative number of conserved (non-universal) sites, and identical bases at these sites, in consensus base sequences of Aspartate and Glutamate family LUCA tRNA species (Davis, 2008a,b). tRNA cognate for amino acids in the Asp family (Fig. 9) jointly conserved 13.4 ± 0.81 (m ± s.e.m.) sites of tRNA-IA^Asn^_3’UUG_ (Fig. 10a) – with universally conserved sites, anticodon, and cofactor/adaptor self-homology excluded. By contrast, only 4.9 ± 0.61 tRNA^Gln^_3’GUU_ sites were jointly conserved with Asp family tRNA. Most jointly conserved tRNA sites retained the same base: 10.4 ± 1.01 in tRNA-IA^Asn^ and 4.1 ± 0.35 in tRNA^Gln^_3’GUU_ (Fig. 10c). The mean number of jointly conserved sites and homologous bases arising between Glu family tRNA species and tRNA-IA^Asn^_3’UUG_, or tRNA^Gln^_3’GUU_, were, respectively, 6.8 ± 2.25 and 4.9 ± 0.61 for sites, and 3.6 ± 0.75 and 2.7 ± 0.67 for bases (Fig. 10b,d). Sites and bases homology exhibited by Asp and Glu family tRNA with tRNA-IA^Asn^_3’UUG_ and tRNA-ID^Gln^_3’GUU_ differed significantly: F_sites_(df 3, 34) = 26.4, *p* = 5.28×10^-9^, and F_bases_(d5f 3, 34) = 17.6, *p* = 4.54×10^-7^ in an analysis of variance, Mean site and base homology were ranked in sequential difference tests as: (tRNA-IA^Asn^_3’UUG_ v. Asp family tRNA) > (tRNA-IA^Asn^_3’UUG_ v. Glu family tRNA) ≈ (tRNA^Gln^_3’GUU_ v. Asp family tRNA) ≈ (tRNA^Gln^_3GUU_ v. Glu family tRNA).

**Fig. 10.**
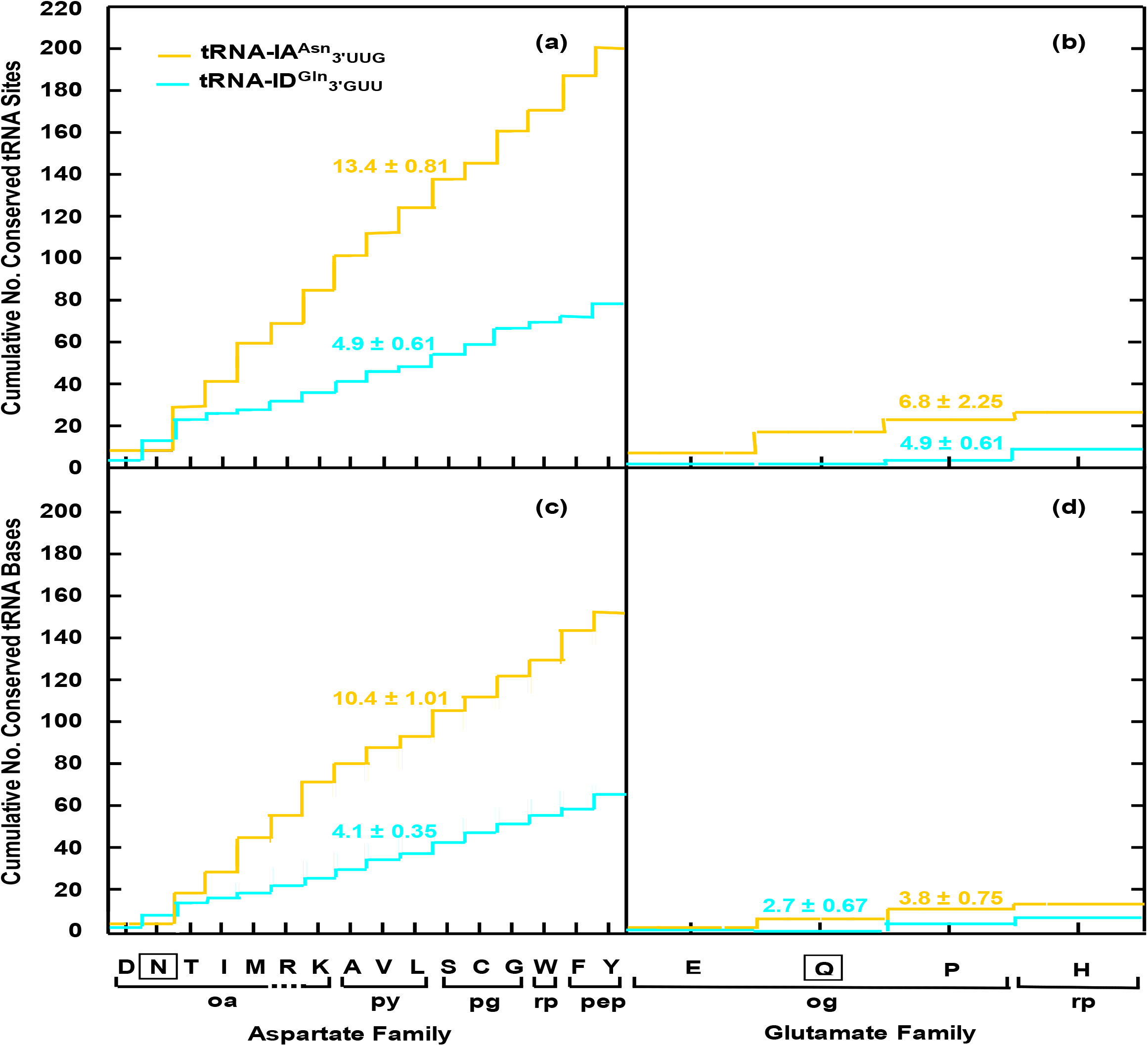
Cumulative number of conserved (non-universal) sites and homologous bases in Aspartate and Glutamate family tRNA species matched with tRNA-IA^Asn^_3’UUG_, or tRNA-D^Gln^_3’GUU_. Single-letter abbreviations, on abscissa, give amino acid specificity of each tRNA; Asn and Gln tRNA (boxed) self-homology excluded. Two-letter abbreviations denote each amino acid precursor, or branchpoint. Mean increments (with standard error) are shown. Site and base homology in Aspartate family tRNA, with tRNA-IA^Asn^_3’UUG_, significantly exceed that of other distributions (see Table S3).

Cofactor/adaptor tRNA of sixteen amino acids with aligned path-distance and depth code evolution metrics (Fig 8a), from four synthesis families with oxaloacetate, pyruvate, phosphoglycerate, or phosphoenolpyruvate as precursor, are revealed by the phylogenetic tree in Fig 9, and corroborated by cumulative sequence homology in Fig. 10, to have diversified from tRNA-IA^Asn^.

## 10 Aspartate family and code duality

Diversification of Aspartate family cofactor/adaptors from tRNA-!A^Asn^ may be seen as evidence that the underlying family of amino acid synthesis pathways diversified from oxaloacetate, *oa.* Central metabolism intermediates upstream of each pathway branch-point retain an invariant, free carboxyl (Fig. 11), allowing attachment of a tRNA cofactor/adaptor and extension of each amino acid synthesis pathway back to *oa* – as the primal precursor (Davis, 2013). Retention of free carboxyl groups by amino acid synthesis intermediates and, by extension, free invariants in central metabolism pathways, were revealed to be the conserved vestige of a cofactor, or scaffold, attachment site (Davis, 2013, 2015, 2018). Since central metabolism reactions predate translation, their role in bridging the gap between *oa* and amino acid synthesis path branch-point, leaves the path-distance metric unaltered; take-over of a Val^5^ path reaction segment, in the synthesis of Ile^7^, exemplifies this mechanism in amino acid synthesis (Davis, 1999).

Arranging Aspartate family amino acids by synthesis branch-point and path-length (Table 2), reveals *oa* as direct precursor to seven amino acids. It contributed amino acids at each stage of code evolution, involving successive recruitment of codon 5’-, mid-, and 3’-site bases (Fig. 1). Beyond initial NH_4_^+^ fixing reactions resulting in conversion of *oa* to Asp^1^, and its amide Asn^2^, and extensions of this pathway, the Aspartate family network of amino synthesis pathways is seen to have expanded, by utilizing *pyr* and *3-pg,* as branch-points in the tRNA-dependent synthesis of six expansion-phase amino acids, with path-lengths of 2 to 8 steps. They entered the code during its expansion, when codon mid-site bases were recruited to encode incoming amino acids. In the final stage of code evolution, the 3’-base in weak-bonding, preassigned codons, was recruited to encode latecomer amino acids. Phosphoenolpyruvate, *pep,* was branchpoint in the long pathways yielding three large aromatic amino acids, Phe^11^, Tyr^11^, and Trp^14^. Addition of Lys^10^, synthesized on a path originating at *oa*, and capture of Arg^9^, completed the Aspartate family.

**Table 2.** Souce of Aspartate Family amino acid precursors at indicated stagesof codon base recruitment

| source | RCC | CT | RPC | CT |
| --- | --- | --- | --- | --- |
| precursor<sup>1</sup> | *oa* | *pyr* | *3-pg* | *pep* |
| | $\begin{array}{c} \text{COOH} \\ \vert \\ \text{CH}_2 \\ \vert \\ \text{C=O} \\ \vert \\ \text{COOH} \end{array}$ | $\begin{array}{c} \text{CH}_3 \\ \vert \\ \text{C=O} \\ \vert \\ \text{COOH} \end{array}$ | $\begin{array}{c} \text{HC-O-P} \\ \vert \\ \text{HC-OH} \\ \vert \\ \text{COOH} \end{array}$ | $\begin{array}{c} \text{CH}_2 \\ \vert \vert \\ \text{C-O-P} \\ \vert \\ \text{COOH} \end{array}$ |
| invariant | COOH | COOH | COOH | COOH |
| tRNA specificity and codon base stage | Asp<sup>1</sup><br>Asn<sup>2</sup> 5'-<br>Thr<sup>6</sup><br>Ile<sup>7</sup><br>Met<sup>7</sup><br>(Arg<sup>9</sup>)<sup>2</sup><br>Lys<sup>10</sup> | Ala<sup>2</sup><br>Val<sup>5</sup><br>Leu<sup>8</sup> | Ser<sup>4</sup><br>Cys<sup>5</sup><br>Gly<sup>5</sup> mid- | Phe<sup>11</sup><br>Tyr<sup>11</sup><br>Trp<sup>14</sup> 3'- |

<sup>1</sup> Arrows indicate direction of precursor recruitment, based on amino acid path-lengths.
<sup>2</sup> Arg<sup>9</sup> acquired an Asn-related cofactor/adaptor in a tRNA exchange (Davis, 2013).

Two unequal subsets of amino acids have been identified within the standard set, exhibiting aligned and misaligned path-distance and phylogenetic-depth metrics (Fig. 8), cofactor/adaptor tRNA sequence homology (Figs. 9, 10), and different primal precursor in each network of tRNA-dependent synthesis pathways (Fig. 1, Table 2). NH_4_^+^ fixer diacid/amide amino acid pairs, Asp^1^/Asn^2^ and Glu^1^/Gln^2^, are at the base of this split. Aminoacylation of their tRNA species is, notably, pair-specific with respect to synthetase class; aaRS-I for Glu^1^ and Gln^2^, and aaRS-II for Asp^1^ and Asn^2^. As synthetases are the product of code-directed translation, with comparatively late-stage residue motifs (Davis, 2008a,b), their distribution among the NH_4_^+^ fixers provides compelling evidence of a duality among ribozymal synthetases at the code origin.

## 11. Discussion

The correlation between amino acid path-distance and phylogenetic-depth (Fig. 8a) validates both as metrics of code evolution. Restriction of the correlation to sixteen amino acids derived from oxaloacetate, pyruvate, phosphoglycerate, or phosphoenolpyruvate, with misalignment versus oxoglutarate derived (Glutamate family, His^13^ included) amino acids (Fig. 8b), furthermore, furnished evidence of a second duality within the code. This raises the possibility it was linked, in some manner, with the family-specific synthetase duality exhibited by NH_4_^+^ amino acids, noted in Sec. 10. Phylogenetic analysis of cofactor/adaptor tRNA sequences (Fig. 9), after elimination of variations accrued in the long interval following species divergence (Eigen et al., 1989), revealed that tRNA, cognate for amino acids with aligned metrics, depicted in Fig. 8a, had diversified from the asparagine cofactor/adaptor. Elevated site, and base, homology with tRNA-IA^Asn^_3’UUG_, exhibited by these tRNA (Fig. 10) affirmed the antiquity of the asparagine cofactor/adaptor. Non-Glutamate family amino acids, thus, formed an extended family. The continuity of advancement in code formation, apparent from the linear relation between amino acid synthesis path length and inferred residue frequency among pre-LUCA proteins (Fig. 8a), implies that sub-families of the 16 member Aspartate family (Supplement, Fig. S1), whose extant synthesis pathways branch from central metabolism at pyruvate, phosphoglycerate, or phosphoenolpyruvate, originated from a single source. With Asp^1^ and Asn^2^ placed as first generation members of the Aspartate family, by path-distance and depth metrics, oxaloacetate arises as the primal precursor to this family of amino acids. Loss of the segment of central metabolism reactions, upstream from current branch-points to oxaloacetate (Table 2), served to eliminate duplicate reaction segments upon the takeover of amino acid synthesis pathways by proteins.

Reconciling path-distance and depth estimates of the time-order of encoding Glutamate family amino acids (Fig. 8) requires identifying the source of a lower than expected level of Glu^1^ and Gln^2^ incorporation into pre-LUCA proteins and a far higher incorporation of the long-path, ring-bearing amino acid, His^13^. Glu^1^ and its amide contribute substantially, as donors in transamination reactions during amino acid synthesis (Michal, 1999). This diverted both of these first generation additions to the code, synthesized on paths of minimal length, from incorporation into the residue sequences of pre-divergence proteins (Fig 8b; Table S2), consistent with lowering their depth index (Brooks et al., 2002).

With a depth index, p_LUCA_/p_mod_, of 1.67 ± 0.17 (mid-range ± half-spread), His^13^ exhibits the steepest rise in residue frequency on retrodicting to pre-divergence proteins (Table S2), while paradoxically having a long, 13-step synthesis (Figs. 1, S1). An additional incongruency arises with it being encoded by a doublet, CAR, within a codon box, which also coding for an NH_4_^+^ fixer, early-comer, Gln^2^. Its long synthesis path and split-box codons indicate that the elevated pre-LUCA incorporation of His^13^, inferred from its depth-index (Brooks et al., 2002), does not result from an early inclusion in the code. With a catalytic potential of 189 (Guttridge and Thornton, 2005; Davis, 2013), far exceeding other amino acids, its appearance, late in code formation, plausibly led to a pre-LUCA surge in the number of His^13^-bearing ‘acid-base’ enzymes. A resulting rise in His^13^ incorporation would directly elevate the depth index of this code late-comer. In this event, His^13^ differs from other late-comers, whose phylogenetic depth correlates with the time-order of amino acid addition to the code, based on their path-distance synthetic order (Fig. 8) and LUCA cofactor/adaptor tRNA homology (Fig. 9).

With oxaloacetate as primal precursor to the 16 amino acids of an extended Aspartate family, and Glutamate family NH_4_^+^ fixers, Glu^1^ and Gln^2^, serving as donors in transamination reactions during amino acid synthesis, a family-based, pre-synthetase duality existed at the origin of the code. It appears likely, therefore, to have been the source of the initial amino acid family-specific distribution of class-I and -II aminoacyl-tRNA synthetases (Sec. 10). Retention of a ribozymal peptidyl-transferase within the large ribosomal subunit (Noller et al., 1992; Agmon et al. 2005; Davidovitch et al., 2009; Petrov et al., 2014) provides compelling support for an era reliant on RNA replicators, catalysts, and cofactors (Orgel, 1986; Gilbert, 1986) in the evolution of translation, including that preceding both classes of aminoacyl-tRNA synthetases. Ribozyme active sites even conserve evidence of pre-RNA stages of life (Davis, 2015, 2018). Analysis of ribosome structure identifies peptide synthesis as a first step in its evolution, leading ultimately to protein synthesis through decoding an ordered base sequence in template-directed translation, read by charged tRNA, equipped (pre-template) with an amino acid acceptor stem and, subsequently, combined acceptor and anticodon stems (Saks and Sampson, 1995).

Codon bias detected in pre-divergence genes indicated assignment of triplet 4-set, UCN, to Ser^4^ preceded allocation of doublet AGY (Diaz-Lazcoz et al., 1995). Conserved 4-set codons of Ser^4^ also significantly exceeded conserved 2-set codons (conserved/non-conserved frequency index > 1.0) in the results reported for six species by Brooks and Fresco (2003, Table 4), with χ^2^_df=1_ 3.87, p = 4.91×10^-2^, notwithstanding the *Aeropyrum pernis* (an archaean) preference for 2-set Ser^4^ codons: number of conserved/non-conserved codons 4-set, 14/10, and 2-set, 2/10. Present evidence reveals initial assignment of UCN triplets to Ser^4^ contributed to saturating the NCN column (Fig. 7), minimizing the risk of mutations to unassigned, untranslatable triplets, during expansion from the NH_4_^+^ Fixers code, at NAN column (Sec. 6). Assignment of NCN column triplets prior to NGN triplets, conforms with selection for codon bonding strength during code evolution (Grosjean and Westhof, 2016); intact UCN box triplets have an estimated bonding enthalpy of −16 kcal/mol versus −14 kcal/mol for split-box AG^Y^_R_ box triplets (Fig. 5a).

With the first code located at the NAN triplet column, it had a mean codon bonding enthalpy estimated at −12.5 kcal/mol (Fig. 1). Back-tracking from its invariant mid-A indicates code formation was preceded by translation on a poly(A) template, with a ratchet-equipped ribosome and tRNA-ID adaptor bearing Asp^1^, or Glu^1^. In this event, the initial stages of code formation were shielded from inter-codon competition, particularly since non-NAN triplets were untranslatable. Vertical (5’-site coding) intra-NAN column diacid/amide amino acid domains, in the NH_4_^+^ Fixers code, reduced the risk of mutation to an untranslatable triplet from 3/4 (random array) to 1/3 (restriction to mid-base substitutions). After allowance for history and ‘lethal’-error avoidance (a result of starting small), selection for codon bonding strength represented a significant determinant of code formation, as Grosjean and Westhof (2016) proposed.

An analysis of codon-bias expanded beyond ordering-in-time the doublet and quadruplet codon set of a single amino acid, to whole rows of the code, indicated that recruitment of GNN codons by Asp^1^/Glu^1^, Ala^2^, Gly^5^, and Val^5^ preceded ANN codon acquisition by Asn^2^/Lys^10^, Thr^6^, Ser^4^/Arg^9^, Ile^7^/Met^7^ (Brooks and Fresco, 2003). Additional to the horizontal direction of time in code evolution, reflecting mid-base substitutions linked to amino acid synthetic order (Fig. 1), this development introduces a vertical component (5’: G ➔ A) of time in code formation. The former is most apparent among related amino acids in both rows: Ala^2^ GC • ➔ Val^5^ GU • (pyruvate derived) and Asn^2^ AA• ➔ Thr^6^ AC• ➔ Met^7^/Ile^7^ AU• (oxaloacetatederived). One of four GNN code boxes is split, Asp^1^/Glu^1^, and path-distance invariant; whereas, three ANN boxes are split, with Asn^2^/Lys^10^ and Ser^4^/Arg^9^ exhibiting 5-to 8-step differences. They contribute to shifting the mean path-distance metric from 2.8 ± 0.92 steps among GNN row amino acids to 6.4 ± 1.04 steps in the ANN row, providing a source of the vertical time component in code evolution. Path-lengths of ANN row amino acids significantly exceed those of GNN amino acids: p = 1.63 x 10^-2^ in a t-test. When long-path ANN amino acids, in split-boxes, are removed, however, its mean amino acid path length, 4.8 ± 1.11, does not differ significantly from the ANN mean: p = 0.11. The vertical component of time in GNN versus ANN rows vanishes; the proposition that GNN row amino acids entered the code before those in the ANN row solely reflects overprinting of ANN codon boxes, in the final stage of code formation. It is notable that the horizontal time-metric is oriented in the same codon mid-base direction, in both rows: GNN, C ➔ U (Ala^2^➔ Val^5^), and ANN, A ➔ C ➔ U (Asn^2^➔ Thr^6^ ➔ Met^7^). Amino acid hydrophobicity and codon bonding strength (after NAN), likewise, share the horizontal orientation of synthesis path-length (Fig. 1). Mid-base recruitment during expansion from the NH_4_^+^ Fixers Code is seen to have occurred in a compact, columnwise pattern, necessitated by the risk of an unreadable triplet stalling translation.

Back-tracking from the NH_4_^+^ Fixers Code indicates the genetic code arose from translation on a poly(A) template involving a ratchet-equipped ribosome and tRNA-ID cofactor/adaptor, bearing either diacid amino acid, and resulting in random sequence polyanionic polypeptides (Davis, 2006). A polymeric source of N atoms, particularly following residue amidation, resulted, within a primal metabolic system at a ferrogeneous cationic surface (Wächtershäuser, 1992; Dodd et al., 2017). Introduction of the pyrimidine complement of A led to the standard codon:anticodon double-helix (A form) within the ribosome proofing center (Ogle et al., 2001) and opened the way for synthesis of polypeptide chains of optimal length (Davis, 2019). The small first code with codons restricted to 5’-specificity, a mid-A, and degenerate 3’-site (Fig. 1) required only four tRNA species, bearing a generic 3’-XUU anticodon, following the proposed transition from base self-recognition to complementarity within the proofing center (Davis, 2019), and retention of ambiguous Asp^1^ and Glu^1^ incorporation, specified by the GAN codon set. This simple code may be noted to reduce the initial risk of mutation to an unassigned, untranslatable triplet from ¾ (random code) to ⅓ (midbase change to NH_4_^+^ Fixers Code).

Code expansion retained this error-minimizing, compact pattern, by recruiting codons in a columnwise manner, through successive mid-base recruitment, C ➔ G ➔ U, in an order consistent with selection for codon bonding strength (Figs. 5, 6; Grosjean and Westhof, 2016). Synthetic order based on path-distance (Fig. 2), broadly equivalent to phylogenetic-depth (Fig. 8), determined the order of amino acid addition to the code. From the four polar amino acids and Ter signal of the NH_4_^+^ Fixers Code, with 16 NAN set triplets (1 code column), ten increasingly hydrophobic amino acids were added to the code during expansion (Fig. 2), a total of 60 triplets assigned (4 code columns, less codon box CGN).

Pre-LUCA proteins furnish an attractor driving the encoding and incorporation of increasingly hydrophobic amino acids during code expansion. A 23-residue antecedent of low potential ferredoxin, pro-Fd-[5] (Davis, 2002), reconstructed by Nørgaard (2009) and Nørgaard et al. (2009), with a code stage 5.6 ± 0.4 residue profile, combined a negatively charged 6-residue ‘foot’ (mean transfer free energy 4.9 ± 2.39 kcal/mol, values from Tolstrup et al., 1994) with a non-polar, 17-residue [4Fe-4S]^2+/+^ cofactor-binding segment (mean transfer energy, −0.83 ± 0.71 kcal/mol, see Supplement, Fig S2). Its structure conforms with one of a protein adaptor, designed to anchor a bound cofactor to a cationic mineral surface. As the Fajan-Paneth principle stipulates, the hydrophobic cofactor binding segment would lengthen the interval of attachment to a cationic mineral surface by this small, ancient metallo-protein, enhancing its [4Fe-4S]^2+/+^ cofactor participation in a primal metabolism, as formulated by Wächtershäuser (1992). This mid-expansion phase protein amplifies the evident anchoring of N atoms by pre-code, random sequence, polyanionic poly(Asp^1^, Glu^1^), suggesting other encoded proteins, likewise, served as cofactor anchors, thereby aiding cofactor participation in an early form of metabolism. Pre-LUCA H^+^ ATPase proteolipid h1 subunit has a code age of 7.0 ± 0.83, indicating membrane encapsulated cells had evolved by the final stages of code expansion. The highly hydrophobic residues of this transmembrane protein have a mean transfer free energy of −1.9 ± 0.25 kcal/mol.

Six of eight code split-boxes occur within the hydropathy clusters centered on NAN and NUN triplets. At the completion of code expansion, triplets in these columns, respectively, encoded four polar NH_4_^+^ fixers as first generation amino acids (Asp^1^, Glu^1^, Asn^2^, Gln^2^), plus a Ter signal, and four non-polar amino acids acquired in the final stage of code expansion (Val^5^, Ile^7^, Met^7^, Leu^8^). Addition of six long-path amino acids, bearing basic and cyclic side chains followed. This entailed codon reassignment, usually as 3’-pyrimidine/ 3’-purine doublets, in codon boxes with sub-threshold bonding enthalpy, > −16 kcal/mol (Fig. 5a). All four codon boxes of the NAN triplet cluster were split by polar amino acids, resulting in (Asp^1^, Glu^1^), (Asn^2^, Lys^10^), (Gln^2^, His^13^) and (Ter, Tyr^11^) polar pairs. Both NUN triplet boxes were split by non-polar amino acids, yielding (Leu^8^, Phe^11^) and (Met^7^, Ile^7^) non-polar pairs. Three of four post-expansion polar amino acids were thus acquired by polar NAN cluster and two of three non-polar amino acids were acquired by the non-polar NUN cluster to yield a homology distribution with a probability of 5.4 x 10^-3^ for random formation (Fig. 3b).

Far higher probabilities have been attributed to the amino acid distribution within the standard code (Gilis et al. 2001; Goodarzi et al., 2004), when evaluating the code as if it was the product of a ‘frozen accident’. From the present perspective, the code arose following evolution of translation with a ratchet equipped ribosome, tRNA-ID cofactor/adaptor, and poly(A) template. It grew incrementally from a small number of amino acids and codons through the interplay of diverse factors, including formation of amino acid synthesis pathways from precursors in central metabolism, diversification of cofactor/adaptor tRNA and polypeptide/ protein functions, error-avoidance, translation fidelity, and amino acid homology.

## Acknowledgement

Prof Eric Westhof drew my attention to advances in the thermodynamics of translation. Generous donations supporting this investigation are also gratefully acknowledged.

## Supplement

### 1. Abbreviations

#### Reaction cycles

**FRC**, formose reaction cycle; **RPC**, reductive pentose-phosphate cycle; **RCC**, reductive citrate cycle.

***Central metabolism:* precursors: oa**, oxaloacetate; **og**, 2-oxoglutarate. **branch points: py**, pyruvate; **pe**, phosphoenol-pyruvate; **gp**, 2-phosphoglycerate; **pg**, 3-phosphoglycerate.

**Amino acids** Standard three-letter and single-letter amino acid designations are used. Lower-case, double-letter abbreviations identify synthesis intermediates and their source (Figs.1, S1).

***Alanine, Ala, A***

***Arginine, Arg, R:* ng**, N-acetyl-glutamate; **np**, N-acetyl-glutamate-phosphate**; ns**, N-acetyl-glutamate-γ-semi-aldehyde**; no**, N-acetyl-ornithine**; or**, ornithine**; cn**, citrulline; **rs**, arginino-succinate.

***Asparagine, Asn, N***

***Aspartate, Asp, D***

***Chain termination, Ter, X***

***Cysteine, Cyst C***

***Glycine, Gly, G***

***Glutamate, Glu, E***

***Glutamine, Gln, Q***

***Histidine, His, H:* pp**, phosphatidyl-ribosyl-pyro-phosphate; **pt**, phospho-ribosyl-adenosine-triphosphate; **rm**, phosphoribosyl-adenosine-mono-phosphate; **ro**, phospho-ribosyl-formimino-amino-imidazole-carboxamideribose-phosphate; **ru**, phospho-ribulosyl-formimino-amino-imidazole-carboxamide-ribose phosphate; **ig**, erythro-imidazole-gylcerol-phosphate; **ia**, imidazole-acetol-phosphate; **hp**, histidinol-phosphate; **ho**, histidinol.

***Isoluecine, Ile, I:* kb**, α-keto-butyrate; **ab**, α-aceto-α-hydroxy-butyrate; **dv**, α,β-dihydroxy-iso-valerate;

***Leucine, Leu, L:* pm**, α-isopropyl-malate; **im**, β-isopropyl-malate; **ic**, α-keto-isocaproate.

***Methionine, Met, M*: sh**, o-succinyl-homoserine; **hc**, homocysteine.

***Lysine, Lys, K:* dl**, α,β-dihydro-picolineate; **pd**, Δ1-piperdiene-2,6-dicarboxylate**; sk**, N-succinyl-ε-keto-α-amino-pimelate**; sa**, N-succinyl-α,ε-diamino-pimelate; **dp**, α,ε-diamino-L-pimelate; **mp**, meso-αε-diamino-pimelate.

***Phenylalanine, Phe, F:* pf**, prephenate; **fp**, phenyl-pyruvate.

***Proline, Pro, P*: gs**, glutamate-γ-semialdehyde; **pc**, Δ1-pyrroline-5’-carboxylate.

**Serine, Ser, S: pp**, phospho-hydroxypyruvate; **ps**, phospho-serine.

***Threonine, Thr, T*: ap**, aspartyl-phosphate; **as**, aspartate-β-semialdehyde; **hs**, homoserine; **ph**, o-phospho-homo-serine.

***Tryptophan, Trp, W*: ah**, β-deoxy-arabino-heptulosonate-7-phosohate; **dq**, 5-dehydroquinate; **ds**, 5-dehydro-shikimate; **sk**, shikimate; **kp**, shikimate-5-phisohate; **es**, 3-enol-pyruvyl-shikimate-5-phosphate; **ca**, chorismate; **aa**, anthranilate; **ra**, N-phospho-ribosyl-anthranilate; **cr**, 1-(o-carboxyphenyl-amino)-1’-deoxyribulose-5-phosphate; **ip**, indole-3-glycerol-phosphate; **in**, indole.

***Tyrosine, Tyr, Y*: hf**, p-hydroxy-phenyl-pyruvate.

***Valine, Val, V:* al**, α-aceto-lactate; **dv**, α,β-dihydroxy-iso-valerate; **kv**, α-keto-isovalerate.

#### Nucleotides

***Adenosine, A; Cytosine, C; Guanosine, G; Uridine, U. Triplet coding site, X.***

### 2. Synthesis pathways in amino acid families

**Fig. S1.**
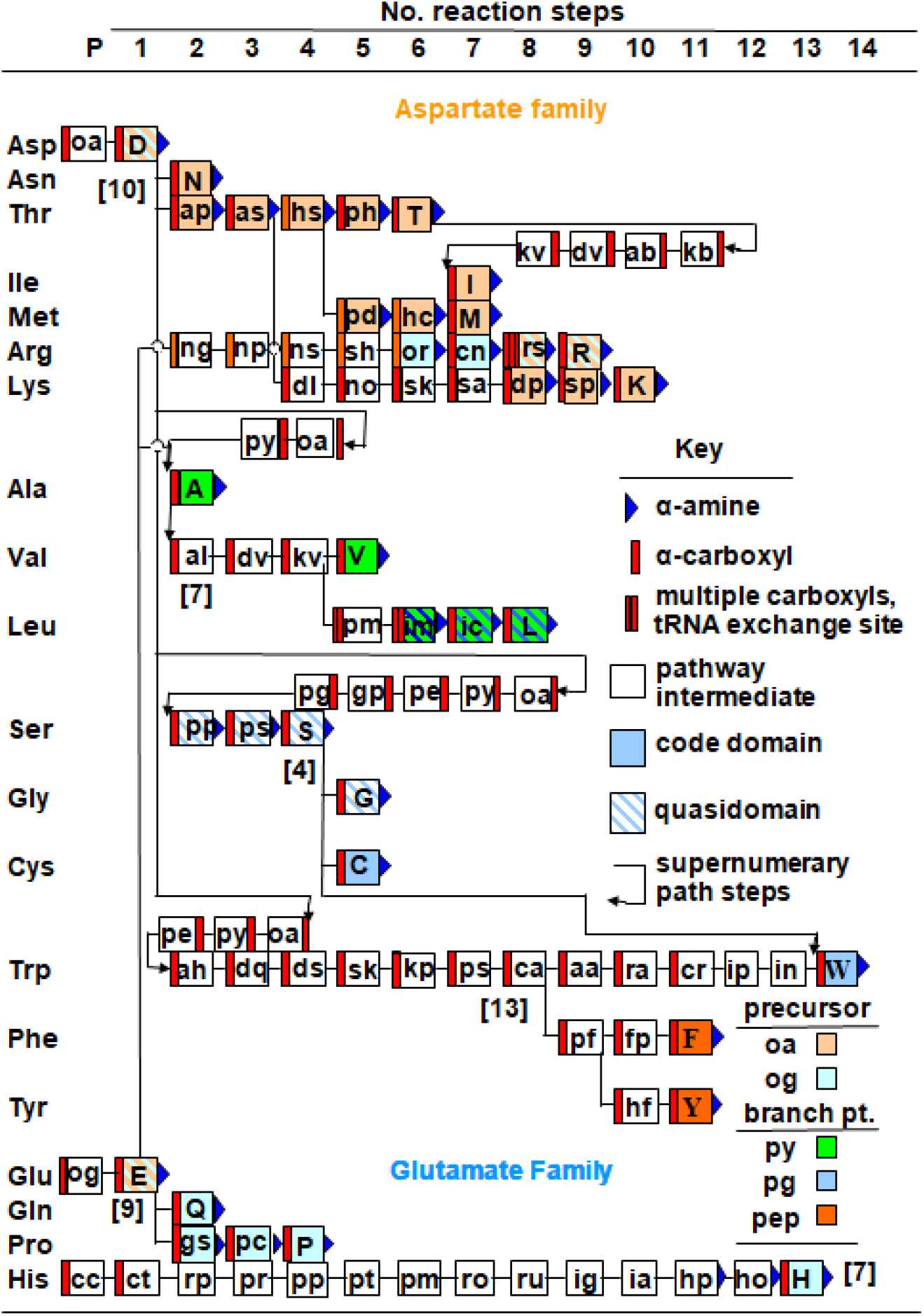
Aspartate and Glutamate family tRNA-dependent amino acid synthesis pathways. Homology with tRNA-IA^Asn^ and carboxylated intermediates upstream from each path branch-point supports oxaloacetate (oa) being primal precursor to four Asp family subfamilies. Small Glu family retained its primal precursor, oxoglutarate. An over-bar gives path-length from precursor, P, with pre-formed RCC and CT steps discounted. Two-letter abbreviations denote amino acid intermediates; red bar signifies a free α-carboxyl (tRNA attachment site), and blue triangle signifies an α-amine. Both families include α-amino acid intermediates, indicative of pathway end-product incorporation. Neighboring codons read by related tRNA, cognate for sibling amino acids, form a domain; quasidomains result on tRNA exchange. Letter and background colors identify precursors and branch-points. The number of path pre-LUCA enzymes is in brackets. Adapted from Davis (2013).

### 3. Hydropathy of protein residues in code expansion phase

**Fig. S2.**
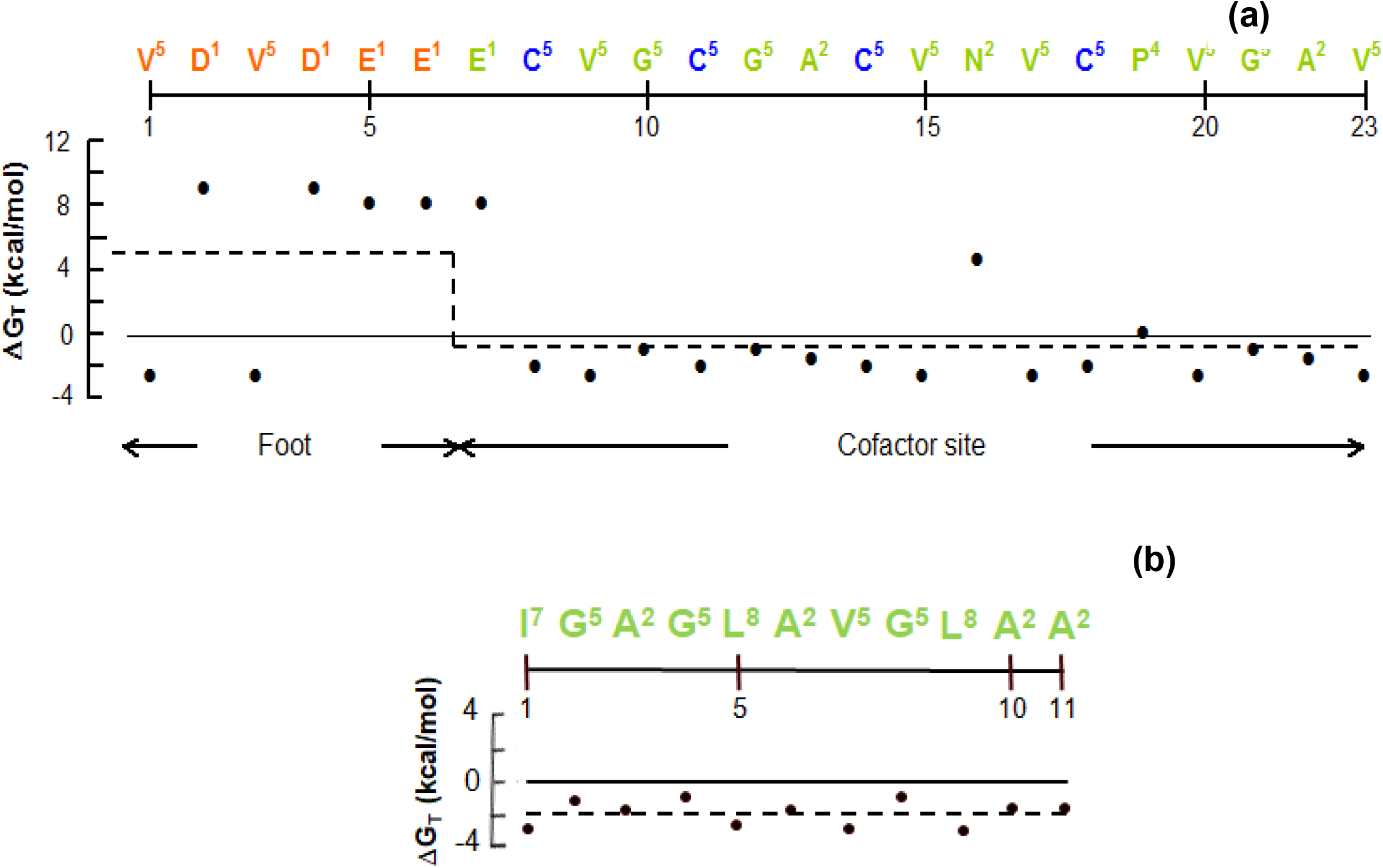
(a). Residue hydropathy in ferredoxin antecedent pro-Fd-[5]. Pre-LUCA low potential ferredoxin antecedent, pro-Fd-[5], contains a negatively charged 6-residue (red), N-terminal foot, with mean hydropathy transfer free energy (ΔG_T_) of 4.9 ± 2.39 kcal/mol (dashed line), connected to a 17-residue C-terminal cofactor binding segment (green) (Davis, 2002; Nørgraard, 2009, Nørgaard et al., 2009). Four Fe atoms coordinately bond to sulfur atoms in cysteine residues (blue) to form a cubic [4Fe-4S]^2+/+^ cluster. With a mean transfer free energy of −0.82 ± 0.71, the hydrophobicity of the long cofactor binding segment would prolong attachment, by Fajan-Paneth’s principle, of this small, ancient metallo-protein to a cationic mineral surface. Amino acid side-chain ΔG_T_ values are from Tolstrup et al. (1994). (b) An 11 residue segment with a late code expansion stage profile identified in the H^+^-ATPase transmembrane proteolipid helix-1 (Davis, 2002). It has a mean transfer free energy of −1.9 ± 0.25 kcal/mol, allowing it to partition with an early cell membrane.

### 4. Hydropathy clusters within the genetic code

**Fig. S3.**
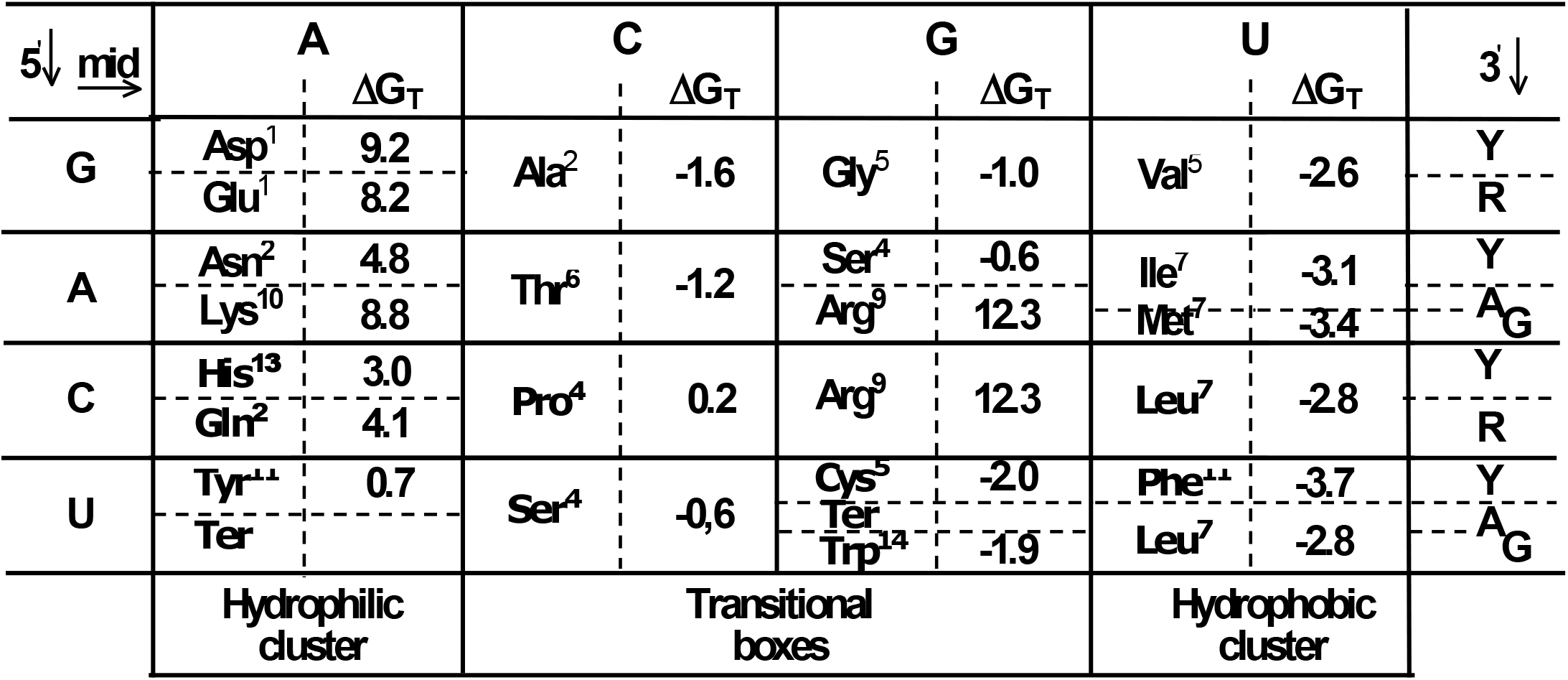
Amino acid transfer free energy distribution within the standard code. Codon mid-base clusters of hydrophilic and hydrophobic amino acids within the standard code, in relation to synthesis path-length (superscripts). Among eight split boxes, five are between a short-path (2 to 7 step paths) and long-path (9 to 14 step paths) amino acid, indicative of pre-assigned codon boxes overprinted by long path amino acids, in the final stage code evolution. Transfer free energy, ΔG_T_, values are from Tolstrup et al. (1994).

### 5. H Bond Enthalpy

**Table S1.**
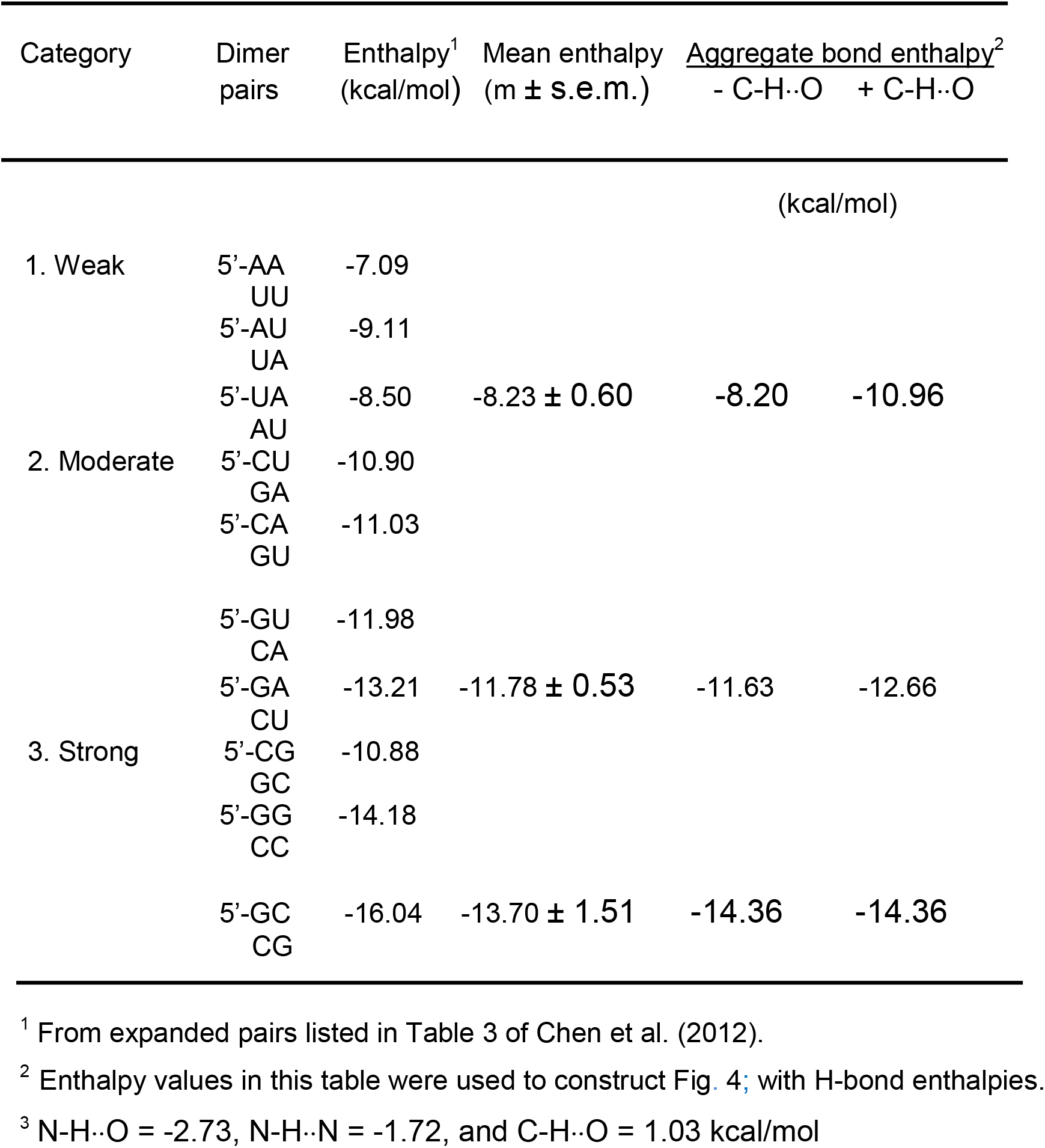
Agreement between dimer pair and inferred H-bond enthalpies

### 6. Time-order of amino acid addition to the genetic code

**Table S2.** Amino acid path distance and phylogenetic depth values^1^

| Amino acid | Path length<sup>2</sup> | Depth $p_{\text{LUCA}}/p_{\text{mod}}$<sup>3</sup> | Category |
| --- | --- | --- | --- |
| Ala | 2 | 1.428±0.070 | 1 |
| Asn | 2 | 1.183±0.113 |  |
| Asp | 1 | 1.064±0.063 |  |
| Gly | 5 | 1.070±0.050 |  |
| His | 13 | 1.670±0.167 |  |
| Ile | 7 | 1.139±0.150 |  |
| Ser | 4 | 1.124±0.112 |  |
| Thr | 6 | 1.282±0.086 |  |
| Val | 5 | 1.200±0.069 |  |
| Met | 7 | 1.105±0.116 | 2 |
| Arg | 9 | 1.037±0.070 |  |
| Pro | 4 | 0.984±0.058 |  |
| Cys | 5 | 0.383±0.077 | 3 |
| Gln | 2 | 0.571±0.089 |  |
| Glu | 1 | 0.914±0.054 |  |
| Leu | 8 | 0.643±0.034 |  |
| Lys | 10 | 0.799±0.049 |  |
| Phe | 11 | 0.590±0.056 |  |
| Trp | 14 | 0.354±0.059 |  |
| Tyr | 11 | 0.464±0.047 |  |

<sup>1</sup> Path-length and depth values that form Fig. 8 correlation.
Glu family amino acids (colored) were treated separately (see text)
<sup>2</sup> See Figs. 1 and S1.
<sup>3</sup> $p_{\text{LUCA}}$ , $p_{\text{mod}}$ are residue frequencies attributed to LUCA and modern proteins by Brooks et al. (2002). Depth values are mid-range ± half-range.

### 7. Amino acid family affiliation of LUCA tRNA species

**Table S3.** Distribution of conserved sites and base identity in LUCA tRNA of aspartate and glutamate amino acid family tRNA versus tRNA-IA^Asn^_3’UUG_ and tRNA-ID^Gln^_3’GUU_

| Amino acid | tRNA-IA<sup>Asn</sup><sub>3'UUG</sub> |  |  |  | tRNA-ID<sup>Gln</sup><sub>3'GUU</sub> |  |  |  |
| --- | --- | --- | --- | --- | --- | --- | --- | --- |
|  | No. sites | Running total | No. bases | Running total | No. sites | Running total | No. bases | Running total |
| *Aspartate family* |  |  |  |  |  |  |  |  |
| D | 9 | 9 | 4 | 4 | 3 | 3 | 3 | 3 |
| N | - | 9 | - | 4 | 10 | 13 | 6 | 9 |
| T | 20 | 29 | 16 | 20 | 10 | 23 | 6 | 15 |
| I | 12 | 41 | 10 | 30 | 3 | 26 | 2 | 17 |
| M | 18 | 59 | 17 | 47 | 3 | 29 | 3 | 20 |
| R | 10 | 69 | 10 | 57 | 3 | 32 | 3 | 23 |
| (3'-UCU) |  |  |  |  |  |  |  |  |
| K | 16 | 85 | 16 | 73 | 4 | 36 | 4 | 27 |
| A | 16 | 101 | 9 | 82 | 6 | 42 | 4 | 31 |
| V | 11 | 112 | 7 | 89 | 4 | 46 | 4 | 35 |
| L | 12 | 124 | 6 | 95 | 2 | 48 | 2 | 37 |
| S | 14 | 138 | 12 | 107 | 6 | 54 | 6 | 43 |
| (3'-AGU) |  |  |  |  |  |  |  |  |
| C | 8 | 146 | 7 | 114 | 5 | 59 | 5 | 48 |
| G | 15 | 161 | 10 | 124 | 7 | 66 | 5 | 53 |
| W | 10 | 171 | 8 | 132 | 4 | 70 | 4 | 57 |
| F | 17 | 188 | 14 | 146 | 3 | 73 | 3 | 60 |
| Y | 13 | 201 | 10 | 156 | 6 | 79 | 6 | 66 |
| Total | **201** |  | **156** |  | **79** |  | **66** |  |
| Mean | **13.4** |  | **10.4** |  | **4.9** |  | **4.1** |  |
| *Glutamate family* |  |  |  |  |  |  |  |  |
| E | 7 | 7 | 3 | 3 | 2 | 2 | 2 | 2 |
| Q | 10 | 17 | 6 | 9 | - | 2 | - | 2 |
| P | 6 | 23 | 3 | 12 | 4 | 6 | 4 | 6 |
| H | 4 | 27 | 3 | 15 | 3 | 9 | 2 | 8 |
| Total | **27** |  | **15** |  | **9** |  | **8** |  |
| Mean | **6.8** |  | **3.8** |  | **3** |  | **2.7** |  |

<sup>1</sup>Number of jointly conserved (non-universal) tRNA sites in LUCA base sequences of cofactor/adaptors for amino acids in Aspartate and Glutamate Families and number of these sites exhibiting base identity with pre-divergence asparagine and glutamine tRNA sequences. Values listed are derived from Davis (2008, Table 2).

## Footnotes

1 Superscripts give amino acid synthesis path-length; one and three-letter amino acid, and one-letter nucleotide abbreviations are used (see Supplement). X, marks the triplet coding site. LUCA, last universal common ancestor.

